# The *Staphylococcus aureus* carotenoid staphyloxanthin modifies the structure of phosphatidylglycerol lipid bilayers

**DOI:** 10.64898/2026.06.25.732829

**Authors:** David Ricardo Figueroa Blanco, Andres Ballesteros, Julian M Delgado, Juan David Orjuela, John Erick Cabrera, Logan Hartmann, Jad Jaber, Kevin Ji, Lucy Knox, Elizabeth Suesca, Gerson-Dirceu López, Chiara Carazzone, Marcela Manrique-Moreno, Gian Pietro Miscione, Stephanie Tristram-Nagle, Chad Leidy, Camilo Aponte-Santamaría

## Abstract

Staphyloxanthin (STX) is a carotenoid synthesized by the human pathogen *Staphylococcus aureus*, providing this opportunistic pathogen with its characteristic golden color. STX protects *Staphylococcus aureus* from oxidative stress by scavenging free radical species. Furthermore, STX has been shown to mechanically strengthen the *Staphylococcus aureus* membrane and to form microdomains that recruit antibiotic-resistance factors, making it an important virulence factor. Thus, inhibition of STX is a promising strategy for intervening against multidrug-resistant strains of this pathogen. The molecular mechanisms by which STX regulates the membrane structure and function of *Staphylococcus aureus* remain unclear. More specifically, the localization of STX within phosphatidylglycerol (PG) bilayers, the primary phospholipid of this bacterium’s membrane, and how this localization drives macroscopic biophysical changes remain unresolved questions. Here, we addressed this issue by integrating molecular dynamics (MD) simulations, X-ray scattering experiments, and fluorescence spectroscopy. We developed an atomistic model of STX, which was validated against X-ray scattering data and which is suitable for all-atom MD simulations. We demonstrate that STX significantly increases lipid packing and acyl chain order of STX–PG bilayer mixtures. In addition, STX self-assembles into clusters, where the long and rigid conjugated triterpenoid chain interdigitates across both leaflets, modifying locally the density of the surrounding PG molecules. These findings provide a molecular explanation to the reduced headgroup spacing and core dynamics observed in fluorescence experiments and are consistent with the formation of structurally-distinct STX-enriched microdomains. Notably, STX reduces the gel-to-liquid crystalline phase transition temperature, indicating a general stabilizing effect for the fluid phase of PG lipids of varying length. Overall, our findings provide molecular insights into how STX enhances the mechanical integrity of PG lipid bilayers and thereby help to establish how membrane remodeling connects with STX’s dual roles, acting as an antioxidant and preventing pore formation.

## Introduction

*Staphylococcus aureus* (*S. aureus*) is a widely spread opportunistic pathogen that inhabits the skin and mucous membranes of approximately a third of the human population (1, 2). *S. aureus* can also survive on various surfaces such as catheters and surgical instruments (3, 4), making this pathogenic microorganism prevalent in hospital-born infections that lead to conditions such as infective endocarditis, pneumonia, soft tissue infections, and which can evolve into life threatening conditions such as bacteremia or sepsis (3). Approximately 20-30% of hospitalized patient bloodstream and surgical site infections have been caused by *S. aureus* (4). This leads to bacteremia incidence ranging from 4.3-38.2 per 100,000 person-years with mortality levels ranging from 10-40% (3, 5, 6). The increased detection of multidrug-resistant *S. aureus* strains (7, 8) and the need to develop alternative treatments drive an effort to understand the structural components of *S. aureus* in more detail.

*S. aureus* synthesizes the lipid carotenoid staphyloxanthin (STX) (9–12). STX is responsible for providing *S. aureus* with its characteristic golden color, and its main function is to protect *S. aureus* from oxidative stress due to the presence of a triterpenoid chain that can sequester free radical species (13). Changes in environmental conditions, such as variations in oxygen levels trigger STX production (14). STX synthesis is controlled through the global stress response regulator SigB (15–17), through two-component systems such as SaeRS (16), MsaB (18), and YjbIH (19). Alternatively, SigB independent regulators such as AirSR trigger STX synthesis in response to increased oxygen levels (20). In general, STX is a stress-response lipid that is closely regulated through molecular sensors that respond to changes in environmental conditions. For these reasons STX is categorized as a virulence factor in *S. aureus* (16, 21), which has led to the exploration of pharmaceutical strategies to inhibit its synthesis (22, 23).

STX content in *S. aureus* varies significantly under different environmental conditions, providing the bacterium with a mechanism to modulate the biophysical properties of its membrane (14, 24). In fact, STX has been found to promote acyl chain ordering, headgroup packing, and membrane bending rigidity in the liquid-crystalline phase of membranes containing phosphatidylglycerol (PG) and cardiolipin (25–28), two of the main lipid components of the *S aureus* membrane in addition to lysyl-phosphatidyl glycerol (29). Consequently, beyond its protective role against oxidation, STX has been tied to an increased resistance to antimicrobial peptides by strengthening the membrane mechanical integrity (25–28).

Carotenoid-enriched regions in *S. aureus* membranes have been shown to form functional microdomains (30). These microdomains recruit a variety of proteins including the penicillin-resistance factor PBP2a, resulting in increased resistance to penicillin. Accordingly, disassembly of these microdomains resulted in increased sensitivity to penicillin in methicillin-resistant *Staphylococcus aureus* (MRSA) strains (31, 32). Sugar functionalized STX molecules were visualized in microdomain fractions from methycillin-resistant strains (31). Moreover, coarse-grained (CG) molecular dynamics (MD) simulations showed the capacity of STX to form STX-enriched structurally-distinct domains (33). Despite all these key observations, the molecular driving factors defining these functional microdomains still remain unclear.

STX features a unique chemical structure (Fig. 1a). STX is a glycosylated 4,4′-Diaponeurosporenoic acid (4,4′-DNPA) bound to fatty acids of varying lengths (10). 4,4′-DNPA is a long 22 carbon triterpenoid attached through an ester bond to the C1 position of glucose. Glucose acts as the polar headgroup of STX. An additional fatty acid is also attached through an ester bond to the C6 glucose position (10, 12). Thus, this specific molecular architecture—characterized by a long, rigid and highly-conjugated chain attached to a shorter acyl chain—is expected to condition membrane properties such as packing, fluidity, and domain formation. CG MD simulations have provided molecular insights into how STX remodels lipid membranes during the formation of lipid domains (33). Nevertheless, a detailed atomistic investigation and experimental validation of the localization of STX in PG lipid bilayers is still missing.

**Figure 1.**
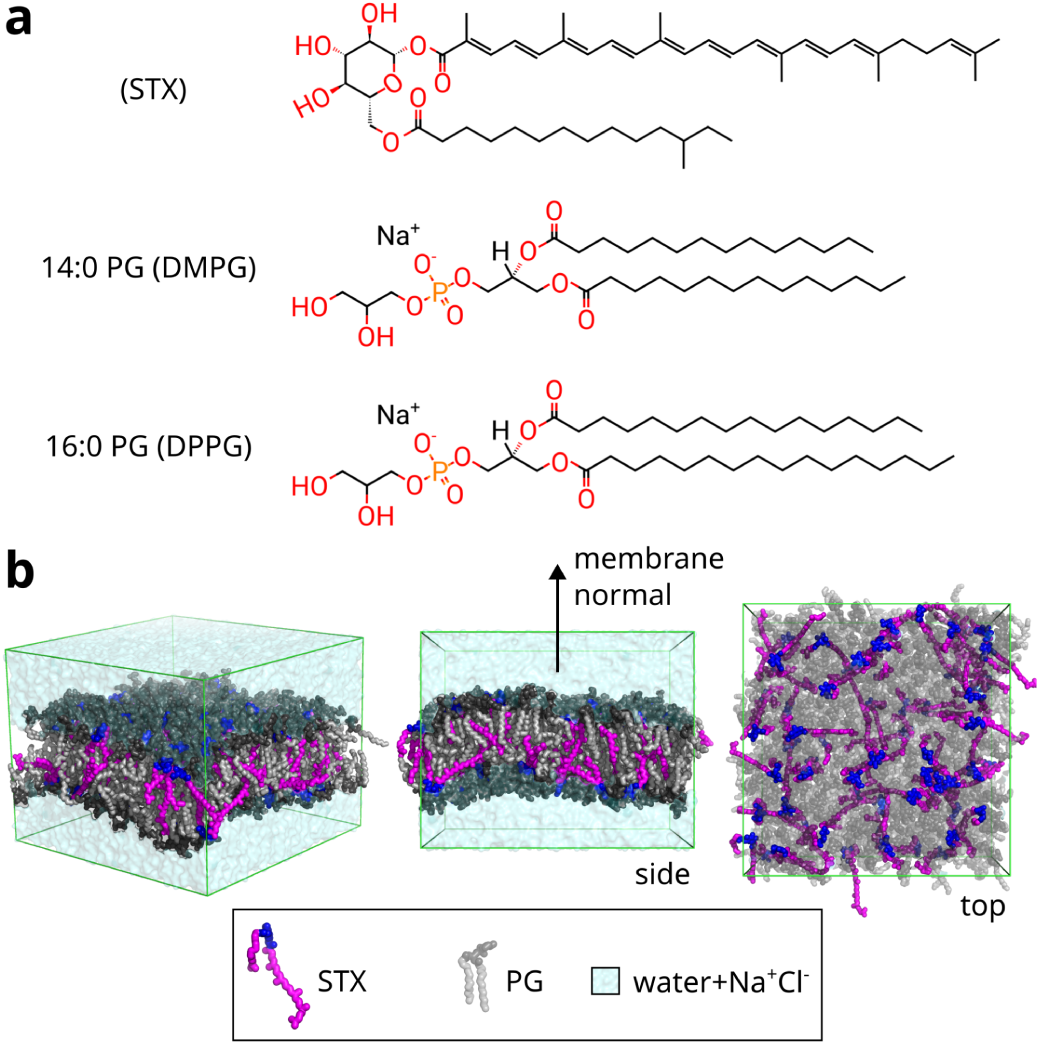
Structure and dynamics of lipid bilayers containing Staphyloxanthin (STX) and phosphatidylglycerol (PG) studied by an integrative approach. **(a)** Chemical structures of Staphyloxanthin (STX), 14:0 PG (DMPG), and 16:0 PG (DPPG) lipids. **(b)** Exemplary configuration of an STX–PG lipid bilayer solvated by explicit water molecules, extracted from MD simulations (here DMPG at a 64:362 STX:DMPG ratio, i.e. 15% mol concentration).

Here, we addressed this question by using minimalistic model membranes containing STX at 0%, 7.5%, and 15% molar ratios and PG lipids, the most abundant headgroup component of *S. aureus* membrane (29), of varying length (DMPG and DPPG). Such simplified models allowed us to systematically investigate the effect of STX on the structure of these bilayers by using MD simulations and X-ray diffuse scattering (XDS) experiments and to contrast our atomistic data with bulk fluorescence spectroscopy measurements, including Laurdan generalized polarization (GP) and DPH anisotropy, and Fourier Transform Infrared (FTIR) spectroscopy.

## Materials and Methods

### Force-field parameter derivation and quantum mechanics (QM) optimization

We generated initial working interaction parameters for STX using the CHARMM General Force Field (CGenFF) (version 2.2.0) (34), by using CHARMM-GUI (35). The O4–C22–C23–C24 dihedral rotation about the C22–C23 bond, joining the ester carbonyl to the conjugated polyene, governs the relative orientation of the saccharolipid headgroup and the rigid chromophore, and is therefore central to the overall structure of STX (Fig. S1a). Thus, we characterized this rotation explicitly by quantum mechanics (QM) calculations. Using Gaussian16 (36), we ran a relaxed DFT dihedral scan over the full 360° in 25 steps of 14.4°, optimizing each conformer at B3LYP/6-31+G(d) geometry, as previously reported in other carotenoid DFT calculations (37), and refining its energy with a single-point MP2/6-31G(d) calculation (MP2/6-31G(d)//B3LYP/6-31+G(d)). We fitted the resulting energy profile to a cosine torsion term. The scan showed a higher rotational barrier than that obtained by the default CGenFF parameters, indicating that this dihedral should be more rigid (Fig. S1a). We therefore increased the height of the cosine terms to match the QM barrier of all dihedral terms involving the C22-C23 bond, and used these corrected parameters throughout all simulations in this study.

### Molecular dynamics (MD) simulations

The simulated lipid bilayers consisted of a mixture of STX and PG lipids at 7.5 or 15% STX mol concentration. For comparison a pure PG lipid bilayer was also simulated. For each of these cases, either DMPG or DPPG lipids were used. See a summary of the simulations in Table 1.

**Table 1.** Summary of the MD simulations.

| [STX] (%)<br>mol) | PG lipid | Temp<br>(°C) | STX:PG No.<br>of lipids<br>ratio (lipids /<br>monolayer) | No. Water<br>molecules<br>per lipid | System<br>size<br>(atom) | No. sim<br>Replicas | Simulation<br>length /<br>replica<br>(ns) | Total<br>Simulation<br>length<br>(ns) |
| --- | --- | --- | --- | --- | --- | --- | --- | --- |
| 0 | DMPG | 45 | 0:128 (64) | 45 | 31642 | 5 | 400 | 2000 |
|  | DPPG | 50 |  |  | 33176 | 5 | 400 | 2000 |
| 7.5 | DMPG | 45 | 32:394<br>(16:197) | ~60 | 124714 | 5 | 400 | 2000 |
|  | DPPG | 50 |  |  | 130095 | 5 | 400 | 2000 |
| 15 | DMPG | 45 | 64:362<br>(32:181) | ~60 | 124714 | 5 | 400 | 2000 |
|  | DPPG | 50 |  |  | 129511 | 5 | 400 | 2000 |

The lipid bilayers were solvated by explicit water molecules. Sodium and chloride ions were added at a 150 mM concentration. In addition, extra sodium ions were added to equalize the negative charge of the PG lipids. The system size ranged from 31642 to 130095 atoms. N=5 simulation replicas were carried out for each system, of 400 ns each, for a total cumulative simulation time of 2 microsecond per system.

The optimized force-field parameters described above were used for STX, while the CHARMM36 force field (38, 39) was employed for the PG lipids. In addition, the CHARMM TIP3P water model (40, 41) and CHARMM ions (42, 43) were used. The systems that only contained PG were generated with CHARMM-GUI (35). The assembly of the systems containing STX was generated as follows. A 2D array of 8×8 STX molecules was generated, by stacking the configuration of a single STX molecule oriented along the z axis. The array was oriented along the xy plane and had dimensions that corresponded to eight times the projected area of a single molecule in each dimension (x or y). In an intercalated fashion, half of the molecules were oriented with the conjugated chain pointing towards negative z values (one leaflet) and the other half towards positive z values (other leaflet). The resulting 8×8 array was used for the simulations at 15 % mol STX concentration. To generate the initial configuration at 7.5 % mol STX concentration, half of the molecules were removed from this array, maintaining them uniformly distributed in space and preserving the same number of STX molecules per leaflet, thus resulting in a system containing 32 STX molecules (16 per leaflet). The resulting arrays were solvated by a preequilibrated lipid bilayer of PG lipids, removing those PG lipids that sterically clashed with the STX molecules, while preserving the desired composition (7.5 or 15 % mol STX) and leaflet symmetry. In practice, 362 PG lipids were considered at 7.5 % mol STX and 394 PG lipids at 15 % mol of STX. To remove voids between the lipids, the solvated array was compressed uniformly laterally in both x and y directions, at a speed of ∼1.8 nm/ps, such that at the end of the deformation the box dimensions along x and y coordinates were approximately 11.8 nm (value estimated as (426 lipids / 128 lipids)^0.5^ x 6.5 nm and taking into account that 128 lipids occupy a box of roughly 6.5൷6.5 nm^2^). Subsequently, the resulting STX–PG bilayers were solvated by explicit water molecules. 0.15 M of sodium and chloride ions were added to the solvent. Moreover, extra sodium ions were added to neutralize the negative charge of the PG lipids. The resulting systems were further equilibrated as described below.

The systems were energy minimized using the steepest descent method and thermalized in the NVT ensemble. Subsequently, they were equilibrated under NPT conditions by imposing position and dihedral restraints on selected atoms of the lipid head groups. Such restraints were gradually removed in six steps. Strength of the restraints and duration of each of the six steps were taken from the CHARMM-GUI equilibration protocol. Production runs followed this gradual equilibration scheme, differing only in one step in which additional deformation was applied to further reduce remaining lateral voids between STX and PG molecules.

The temperature was kept constant (at 45 °C for bilayers containing DMPG and 50 °C for bilayers containing DPPG) by coupling separately the lipid bilayer and the water and ions to the Nosé-Hoover thermostat (44) (coupling time of 1 ps). The Parrinello-Rahman (45) barostat was used to maintain the pressure constant at a value of 1 bar, semi-isotropically along the lipid bilayer plane and perpendicular to it, and with a coupling time constant of 5 ps. Both thermostat and barostat corrections were applied every 20 integration steps.

Short range non-bonded interactions were modeled through a Lennard Jones potential while electrostatic interactions were treated with the particle mesh Ewald method (46, 47). Neighbour atoms for non-bonded interactions were updated according to the Verlet Buffer scheme (48). For the lipid molecules, bonds to hydrogen atoms were constrained with the Lincs algorithm (49). For the water molecules, both bonds and angles were constrained using Settle (50). Equations of motion were numerically integrated, by using the Leap Frog algorithm, at discrete time steps of 2 fs. The molecular dynamics simulation package GROMACS (version 2025.2) was used (51).

### Simulation analysis

#### Electron density profiles

Atomic electron density profiles (EDPs), ⍴(z), were computed for each lipid atom type separately along the coordinate orthogonal to the membrane plane, by using MDanalysis (52). For the simulations including STX the undulation profile was corrected (53, 54). Because of their smaller size, such correction was not necessary for the pure PG bilayers. The profiles were generated at a bin resolution of 0.01 nm, symmetrized with respect to the bilayer center, and stored in *sim* format for further usage to compute the form factors. Atomic density profiles were grouped into the different lipid moieties and presented as a function of the position *z* from the bilayer center. Density profiles were also computed for the conjugated chains of STX, separately for each lipid leaflet, to examine their level of interdigitation.

#### Membrane thickness

The following estimates of the membrane thickness were obtained from the resulting electron density profiles (55–58). D_HH_ was computed as the distance between total density maxima across the bilayer center, after applying a Gaussian smoothing (sigma=0.05 nm). D_B_ was estimated as the Gibbs dividing surface of the water density and 2D_C_ (the hydrocarbon thickness) was defined as the Gibbs dividing surface of the hydrocarbon density. In the two latter cases, the interface region was defined as the region [z_l_ , z_r_] in which the density increased from a value ⍴_l_ to a value ⍴_r_. Accordingly, the position of the Gibbs dividing surface, denoted here as z_G_, was obtained from the equality

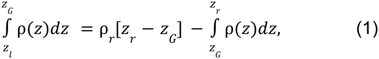

which can be simplified into

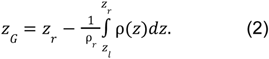

Accordingly, D_B_ and 2D_C_ were estimated as 2z_G_, plugging the water density and the hydrocarbon density, respectively, into eq. (2). For D_B_, the upper boundary, z_r_, was chosen at the position where the density surpassed a value of ⍴_r_ = <⍴_bulk_> -3σ(⍴_bulk_), with ⍴_bulk_ the bulk water density, and the lower boundary, z_l_, at the position where the water density reduced to ⍴_l_ =⍴_r_ /100. Analogously, for 2D_C_, the lower boundary, z_l_, was chosen at the position where the density reached a value of ⍴_r_ = 0.99<⍴_HC_>, with ⍴_HC_ the hydrocarbon density, and the upper boundary, z_r_, at the position where the hydrocarbon density reduced to ⍴_r_ =⍴_l_ /100. The integral in eq. (2) was evaluated numerically using the Simpon method. Note that the thickness estimates derived here (from the density profiles) are equivalent to those that would be derived from the volume probabilities, because densities and volume probabilities are related by a constant volume factor (59).

#### Form factors

For comparison with the X-ray experiments (see below), form factors were derived from the simulations, by taking the Fourier transform of the electron density distributions using the SIMtoEXP package (see below) (60).

#### Local Membrane thickness

The membrane thickness was also monitored locally, using g_lomempro (61). In brief, The distance across the bilayer center between selected headgroup atoms was computed, separately for each lipid. The O7 atom of the glucose group of STX and the phosphorous atom of the phosphate group of the PG lipids were considered for this distance calculation.

#### Area per lipid

The area per lipid was computed using g_lomempro (61). In brief, the area of each lipid was computed as the lateral Voronoi area occupied by it, considering the O7 atom of STX and the phosphorous atom of the PG lipids as reference.

The expectation values of both the local membrane thickness and the area per lipid were obtained at each time frame by averaging over all lipids. These two properties were also studied separately for STX and PG, by selecting at every time frame a random lipid (either STX or PG). Uncorrelated data samples were taken at time frames spaced by the correlation time. Accordingly, the correlation time was estimated as the time when the autocorrelation function decayed to zero. Correlation times were obtained from the resulting time series for either local membrane thickness or area per lipid, by using the *gmx analyze* GMX tool (62).

For the PG lipids, the local membrane thickness and area per lipid were also monitored as a function of their nearest STX lipid molecule.

#### Order parameters

The deuterium-order parameter, S_CD_, of the i−th carbon atom of the acyl chains (C_i_) of the PG lipids was calculated using the following formula (63): S_CD_ = 2S_xx_/3+S_yy_/3, where S_xx_ and S_yy_ are defined as S_xx_=<3cos^2^β-1>/2 and S_yy_=<3cos^2^ɑ-1>/2. Here, <> denotes ensemble average. β is the angle between the vector normal to the membrane plane (i.e. z^^^) and the vector normal to the plane defined by C_i−1_, C_i_ and C_i+1_. α is the angle between z^^^ and the vector in the plane defined by C_i−1_, C_i_ and C_i+1_ which is perpendicular to the vector connecting C_i−1_ to C_i+1_. Order parameters were obtained as time-averages over the concatenated MD trajectories using the GROMACS *gmx order* analysis tool.

#### STX conjugated chain orientation

The orientation of the STX conjugated chain was monitored by computing the angle, θ, corresponding to the inclination of the vector connecting carbons C23 and C41 with respect to the membrane normal, by using the GROMACS *gmx angle* tool.

#### STX contact area

To quantify the level of clustering of the STX lipids, the contact area between them was quantified as the ratio (∑_i_S_i_ - S_T_)/∑_i_S_i_, where ∑_i_S_i_ is the sum of the individual surface areas of the STX molecules and S_T_ is the exposed area of the STX clusters (all STX molecules together). This ratio takes a minimum value of zero, when the STX lipids are not in contact and thus all their surfaces are exposed (∑_i_S_i_ = S_T_), and increases as the STX molecules enter into contact forming aggregates. Accordingly, this quantity relates to the extent of formation of STX clusters. Per lipid and total surface areas were computed with the GROMACS *gmx sasa* tool.

#### Maximum cluster size of STX molecules

The formation of STX clusters was also monitored, by assuming that two STX molecules belonged to the same cluster if they were at a distance smaller than 0.35 nm. The maximum size of all existing clusters at each time was tracked through the GROMACS *gmx clustsize* tool. Subsequently the distribution of the maximum cluster size was determined.

#### Statistical analysis

Unless stated otherwise, the first 150 ns of each simulation replica were accounted as equilibration time and thus discarded from the calculations. Distributions or time-averages over the concatenated trajectories of the quantities of interest were generated. Errors were estimated as the standard error of n=5 simulated replicas or n uncorrelated replicas for the case of the local thickness and area per lipid calculation.

Simulation input, coordinate, topology and parameter files as well as all the scripts to set up and analyze the simulations are accessible at the Edmond data repository here: https://doi.org/10.17617/3.TTF8IJ. In particular, the STX atomic interaction (force-field) parameters, in GROMACS format, can be found there.

#### Molecular visualization

2-dimensional representations of STX and PG lipids were generated with rdkit (64) Visual Molecular Dynamics (VMD) (65) and PyMol (66) were used to visualize the simulated bilayers and to render the simulation snapshots.

### Materials

#### Sample preparation

The synthetic lyophilized lipids 1,2-dimyristoyl-sn-glycero-3-phospho-(1’-rac-glycerol) sodium salt (DMPG) and 1,2-dipalmitoyl-sn-glycero-3-phospho-(1’-rac-glycerol) sodium salt (DPPG) were purchased from Avanti Polar Lipids (Alabaster, AL, USA) and used as received. Staphyloxanthin (STX) was purified from *Staphylococcus aureus,* as described below. The chemical structures of the lipids and STX are presented in Fig. 1a. HPLC-grade organic solvents were purchased from Sigma-Aldrich (St. Louis, MO, USA. Ethylenediaminetetraacetic acid (EDTA) was purchased from Amresco (Solon, OH, USA). Luria–Bertani (LB) medium was prepared with NaCl (ACS, J.T. Baker, USA), tryptone (OXOIO, Basigstoke, Hampshire, UK), and a yeast extract (Dibico, Mexico D.F., Mexico). HPLC-water was obtained from a water purification system: Heal Force Smart-Mini (Shangai, China). The fluorescent probes 6-Dodecanoyl-2-Dimethylaminonaphthalene (LAURDAN) and diphenylhexatriene (DPH) were purchased from ThermoFisher Scientific (Waltham, MA, USA).

#### Purification of staphyloxanthin

The complete purification method has been published previously (33, 72). Briefly, a single colony of the methicillin-susceptible *S. aureus* strain (SA401) was grown in 10 mL of LB medium at 37 °C overnight (16 h) with constant agitation (250 rpm). LB medium contained, per liter, 10 g of NaCl, 10 g of tryptone, and 5 g of yeast extract. Cells were then diluted (1:1000) in fresh LB medium and cultivated for 24 h. Subsequently, the cell pellet was obtained by centrifugation at 7300 x g at 4 °C for 10 min (Thermo Scientific, USA), frozen at −80°C, and lyophilized for 24 h (LABCONCO, Kansas City, MO, USA).

The carotenoid extraction was performed using the previously-reported method (10, 28). One gram of freeze-dried and ground cells were dissolved in 20 mL of methanol containing BHT at 0.1% (w/v); then, 20 glass beads were added and vortex-mixed for 5 min. After that, carotenoids were obtained by centrifugation at 7300 x g for 10 min at 4 °C. The described steps were repeated twice with 10 mL of methanol-BHT (0.1%, w/v) each time. Next, all methanolic phases were combined, and a mixture of ethyl acetate and 1.7 M NaCl (1:3 v/v) was added. The whole mixture was vortexed and then centrifuged at 7300 x g for 15 min at 4 °C. Finally, all the organic phases were evaporated in a rotary evaporator in a water bath at 35 °C. For protein removal, the extract was solubilized in 20 mL of chloroform:methanol mixture (2:1) and transferred to a borosilicate glass tube with a phenolic cap and poly-tetrafluoroethylene (PTFE)-faced rubber liner. Then, 1.7 M NaCl solution was added, and the solution was shaken. Phase separation was achieved by centrifuging at 630 x g for 15 min at 4 °C, and the lower chloroform phase containing carotenoids was collected. Next, the organic phase was dried with anhydrous sodium sulfate, transferred to an amber tube, and finally dried with a CentriVap refrigerated vacuum concentrator (LAB-CONCO, Kansas City, MO, USA).

The separation of staphyloxanthin was performed by applying the carotenoid extract on a preparative thin layer chromatography (PTLC) plate with dimensions of 10 cm height x 20 cm width, the STX purification was made using mobile phase toluene:methanol (84:16) in 25-minutes runs (28). STX was recovered from silica using a mixture of methanol:chloroform (2:1; v/v), dried with anhydrous sodium sulfate, and the solvent was evaporated with a CentriVap refrigerated vacuum concentrator. Subsequently, the identification of the STX was achieved using the liquid chromatography-mass spectrometry (LC-MS) method (10, 28).

### Low-angle X-ray diffuse scattering (LAXS, XDS)

#### Sample Preparation

DMPG and DPPG were mixed with 0, 5, 7.5, 10, and 15 mol% STX (STX/(STX+PG)) in organic solvent, which was then evaporated. Organic solvent (3:1 chloroform:hexafluoroisopropanol) was added to 4 mg of the dry mixture, vortexed, and plated onto Si wafers (15mm W x 30mm L x 1mm H) inside a fume hood while solvent evaporated. Oriented samples consisting of stacks of ∼1800 bilayers were prepared using this “rock and roll” method, where rocking the sample during solvent evaporation creates an immobile, well-oriented film (67). Samples were trimmed to occupy 5mm W x 30mm L along the center of the Si wafer, fixed to a glass block (10mm H x 15mm W x 32mm L) using heat sink compound (Dow Corning, Freeland, MI), and stored in a refrigerator at 4°C. Transfer of samples into an insulated hydration chamber, held at 45°C for DMPG and 50°C for DPPG, immediately prior to x-ray data collection, caused 100% hydration through the vapor within 10 minutes (68). Milli-Q water (MilliporeSigma, Burlington, MA, USA) in the bottom of the hydration chamber kept the samples fully hydrated. After X-ray, thin layer chromatography was carried out to assess the extent of lipid degradation using a solvent phase of 46:18:3 (v:v:v) chloroform: methanol: 7 N NH_4_OH on silica gel 60 plates (EMD, Darmstadt, Germany). Spots were stained with Molybdic acid and iodine vapor. As shown in Fig. S2, only 5% breakdown in DMPG occurred during X-ray at 45 °C while no breakdown of STX occurred.

#### X-ray Data Collection

LAXS data were obtained on two separate trips to station ID7B2 at the Cornell High Energy Synchrotron Source (CHESS, Ithaca, NY) using an X-ray wavelength of 0.8856 Å, beam size 0.26 mm H, 0.35 mm V, and an Eiger 16M detector (Dectris, Baden, Switzerland) distanced 400.1 mm and 348 mm from samples. For each scan the samples rotated either from 0 to 5 degrees during 10 seconds of x-irradiation or from -1 to 6 degrees during 15 seconds of x-irradiation. Background data from air, water, and mylar scattering were collected for 10 or 15 seconds at an x-ray angle of incidence of -2 degrees, where the sample does not contribute to XDS.

#### Data Analysis

The background scattering was subtracted from the sample scattering image to increase the signal-to-noise ratio. Additional background data were removed by interpolating intensity data adjacent to either side of the sample scattering pattern. “Lobes” of diffuse scattering appear due to membrane fluctuations as the sample nears full hydration (69). The lobes are fit using a non-linear least squares analysis to the free energy functional from liquid crystal theory (70, 71),

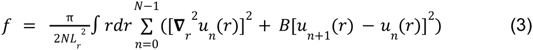

where *N* is the number of bilayers in the stack, *L_r_*is the horizontal (r) domain size, and *K_C_* is the bending modulus. *K_C_*, the elastic bending modulus, directly correlates to the rigidity of a single bilayer, where *u_n_* is the vertical displacement of the nth bilayer and *B*, the compressibility modulus, accounts for interactions with adjacent bilayers in the stack.

The form factor is derived from the intensity data along the entire q_z_ range from 0.2 to 0.7Å^-1^. First, the intensity versus q_z_ is obtained from the liquid crystal theory fit to the XDS data. Then, the Lorentz and absorption corrections are applied when the square root is taken:

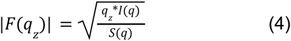

The form factor data are fit through the Fourier transform to a model of a real-space scattering-density profile (SDP) of a lipid bilayer using volume and electron density information (55, 57). Component groups are fit as Gaussians into the total electron density to determine their positions. Only the pure lipids’ EDPs were derived experimentally using SDP, due to the large positional variation of STX component groups within the bilayer, and the lack of volume information about STX.

##### SIMtoEXP

SIMtoEXP performs the Fourier transform of the simulated EDP, which gives the simulated form factors (60). The simulated form factors were compared directly to experimental form factors, thereby avoiding any model assumptions. Since the experimental form factors are obtained on a relative scale, a least-squares analysis was used to scale them to the simulated |F(q_z_)|. Simulated form factors were generated for DMPG and DPPG with 0, 7.5 and 15 mol% STX and compared to experimental form factors.

### Fluorescence Spectroscopy

DMPG and DPPG lipids were dissolved and mixed in chloroform containing 2% vol/vol methanol with varying concentrations of STX incorporated (molar percentages of 5, 10, 15, and 20 mol%). DPH and Laurdan dyes were added to the dispersion at a concentration of 1:150 (probe to lipid molar ratio). The mixtures were dried under a stream of nitrogen and lyophilized overnight to remove residual chloroform. Then, the obtained films were hydrated using 500 µl of HEPES buffer (20 mM HEPES, 190 mM NaCl, 400 mOsm and pH 7.4) at 50 °C to form multilamellar vesicles, to a final concentration of 1.5 mM under periodic agitation on a vortex mixer. In the case of the samples containing DPH, large unilamellar vesicles were formed by extruding the sample 20 times through a 100 nm polycarbonate membrane. For DPH and Laurdan measurements, 34 µl of MLVs or LUVs containing the fluorescent probes were added to a cuvette containing 1 ml of HEPES buffer.

Generalized polarization (GP) and anisotropy (r) measurements were performed in an ISS-PC1 (ISS, Champaign, IL, USA) photon-counting spectrofluorometer equipped with a temperature controller, and sample homogeneity was maintained through continuous magnetic stirring (23). A combination of 0.5-2.0 mm slits was used in the excitation and emission monochromators to set the bandpass to 2, 4, or 8 nm, depending on the sample intensity and photobleaching sensitivity (23). The Laurdan GP value was calculated using the expression:

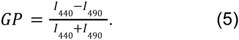

Here, *I_440_*and *I_490_* are the fluorescence intensities at emission wavelengths of 440 nm (gel phase) and 490 nm (liquid-crystalline phase), respectively.

DPH fluorescence anisotropy was calculated according to the definition:

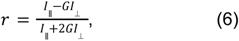

where *I_||_* is the fluorescence intensity when the angle between polarizers is 0° and *I*_⊥_ is the fluorescence intensity when the angle between polarizers is 90° (27), where the G-factor was calculated and the anisotropy corrected automatically by the equipment at each data point. Measurements were performed from 10 to 50 °C (heat rate: 1.5 °C/min). The data presented in the figures represent mean values and standard error of 4 measurements in four independent experiments.

### Infrared spectroscopy experiments

Supported lipid bilayers (SLBs) were prepared in situ in a BioATR II cell. The unit was integrated with a Tensor II spectrometer (Bruker Optics, Ettlingen, Germany) with a liquid nitrogen Mercury Cadmium Telluride (MCT) detector using a spectral resolution better than 0.4 cm^-1^ and 120 scans per spectrum. The required temperature was set by a Huber Ministat 125 computer-controlled circulating water bath (Huber, Offenburg, Germany) with an accuracy of ± 0.01 °C. First, the background was recorded using buffer (20 Mm HEPES, 500 mM NaCl and 1 mM EDTA) in the same temperature range as samples. Stock solutions of pure DMPG and DPPG were prepared in chloroform. Subsequently, the cell was filled with 20 µl of a 20 mM lipid stock solution. Then, STX was added in different concentrations (5, 10, 15, and 20 mol%) in chloroform, and the solvent was evaporated for 20 seconds, resulting in a lipid multilayer film. The cell was filled with 20 µL buffer solution immediately after removal of chloroform for in situ measurements and incubated above the phase transition temperature for 10 min. To determine the position of the vibrational band in the range of the second derivative of the spectra, all the absorbance spectra were cut in the 2970-2820 cm^-1^ range, shifted to a zero baseline, and the peak-picking function included in OPUS 8.8.4 software (Bruker Optics, Ettlingen, Germany) was used. The results were plotted as a function of the temperature. To determine the transition temperature (T_m_) of the lipids, the curve was fitted according to the Boltzmann model to calculate the inflection point of the obtained thermal transition curves using the OriginPro 8.0 software (OriginLab Corporation, Northampton, MA, USA).

## Results

The effect of STX on the structural organization of STX–PG lipid bilayers was studied here by an integrative approach combining MD simulations, X-ray scattering, Laurdan generalized polarization, DPH fluorescence anisotropy, and FTIR spectroscopy.

### Experimental Validation of MD Simulations

The consistency of the generated force field parameters of STX was checked by calculation of the dihedral angles along the conjugated chain. As expected, these dihedrals fluctuated around ±180°, indicating that this chain stayed mostly straight in a highly planar *trans* configuration (Fig. S1b). The STX parameters were validated against X-ray scattering data. Fig. 2 shows the form factors |F(q_z_)| obtained from XDS (Fig. 2a). Their dependency on the STX concentration is analyzed in detail below. To create a model-independent comparison of experimental data to MD simulation, the simulated form factors were also obtained by Fourier transform of the simulated electron density profiles, ρ(z), along the coordinate perpendicular to the membrane z (Fig. 2b). The experimental form factors were scaled to match the amplitude of simulated ones and thus facilitate their comparison (Fig. 2c). Excellent agreement between experimental and simulated |F(q_z_)| was obtained at the minima as well as at the amplitudes of the lobes, not only for pure PG lipid bilayers (Fig. 2c, left) but also for those containing 15% STX (Fig. 2c, right).

**Figure 2.**
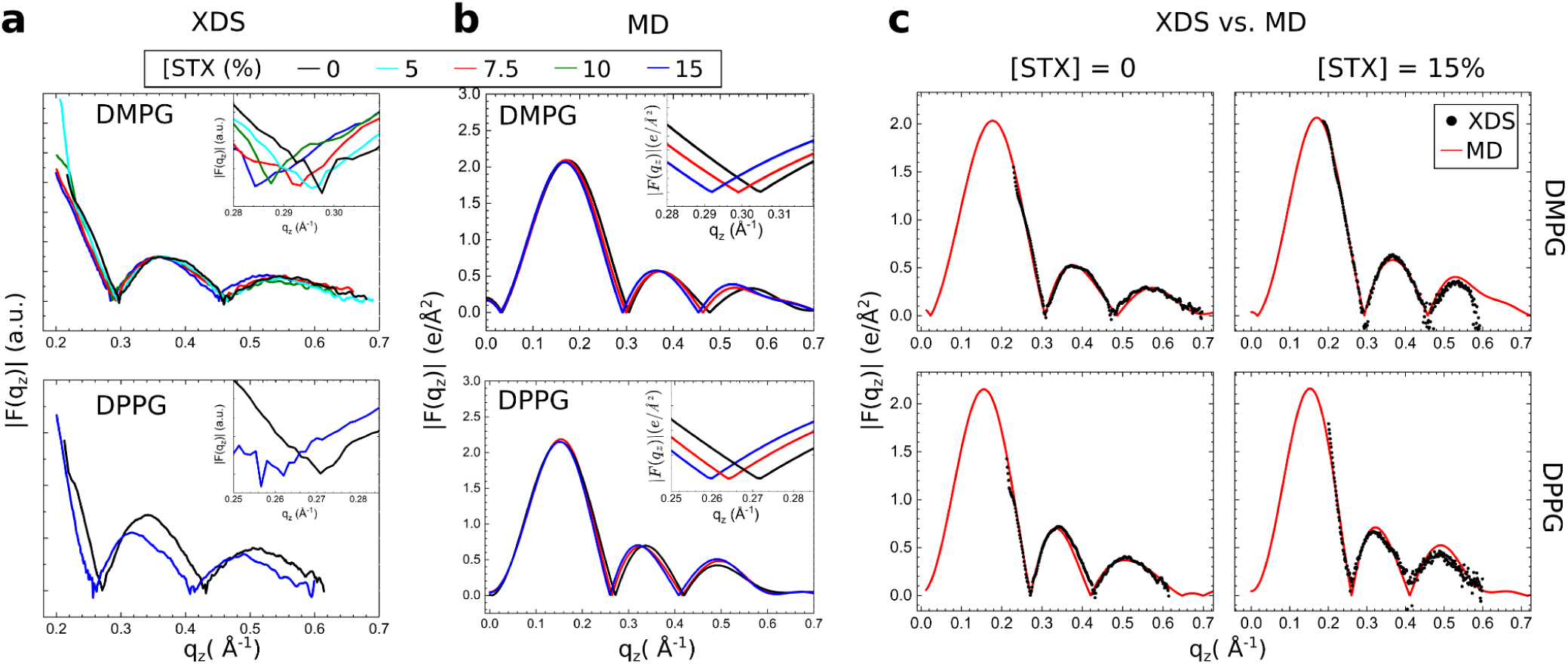
Form factors STX–PG lipid bilayers elucidated by X-ray scattering and MD simulations. Form factors |F(q_z_)| from X-ray diffuse scattering (XDS) experiments **(a)** and from MD simulations **(b)** of DMPG and DPPG at the indicated mol% STX concentrations. The lobes shift to lower q_z_ as STX content increases. **(c)** Comparison of the experimental and simulated form factors for the indicated PG lipid (rows) and STX concentrations (columns). For STX in DPPG bilayers, due to difficulties with the sample preparation at intermediate STX concentrations, XDS was only successful for either pure DPPG bilayers or for those containing 15% mol of STX. Both experiments and simulations were carried out at 45°C for DMPG and 50°C for DPPG.

We also obtained experimental EDPs for pure PG lipid bilayers, via SDP, by using previously published volumes for DMPG (1189 Å^3^ at 45°C) and DPPG (1070 Å^3^ at 50°C) (57) (Fig. S3a,b). These profiles compare well with those extracted from the simulations (Fig. S3c,d). More quantitatively, structural parameters derived from the experimental EDPs were within 3% of those extracted from the simulations, with the exception of the 2Dc which deviated up to 5% (Table 2). These results further confirm the consistency between the experimental and the simulation data for both DMPG and DPPG.

**Table 2.**
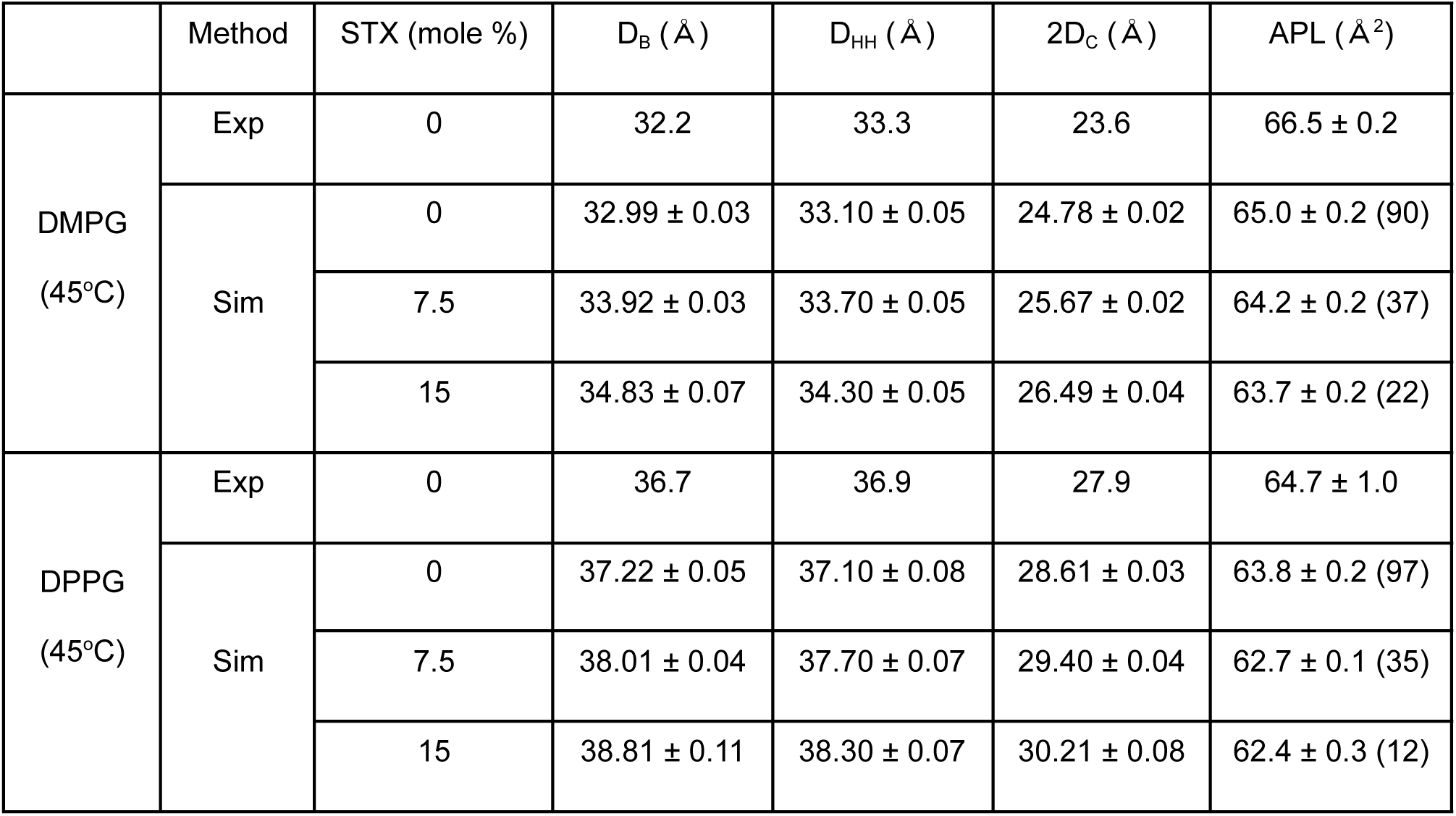
Structural parameters of DMPG (45°C) and DPPG (50°C) lipid bilayers for experiments (0 mol% STX) and MD simulations (0, 7.5, 15 mol% STX). D_B_ (Gibbs dividing surface of the water density), D_HH_ (the distance between total density maxima across the bilayer center), 2D_C_ (hydrocarbon thickness, defined as the Gibbs dividing surface of the hydrocarbon density), and area-per-lipid (APL). Average ± Standard Error of the Mean is shown for the simulations from n=5 independent simulation replicas. The values in parenthesis for the APL indicate the effective number of independent uncorrelated replicas (see methods).

| | Method | STX (mole %) | $D_B$ (Å) | $D_{HH}$ (Å) | $2D_C$ (Å) | APL (Å<sup>2</sup>) |
| --- | --- | --- | --- | --- | --- | --- |
| DMPG<br>(45°C) | Exp | 0 | 32.2 | 33.3 | 23.6 | 66.5 ± 0.2 |
|  | Sim | 0 | 32.99 ± 0.03 | 33.10 ± 0.05 | 24.78 ± 0.02 | 65.0 ± 0.2 (90) |
|  |  | 7.5 | 33.92 ± 0.03 | 33.70 ± 0.05 | 25.67 ± 0.02 | 64.2 ± 0.2 (37) |
|  |  | 15 | 34.83 ± 0.07 | 34.30 ± 0.05 | 26.49 ± 0.04 | 63.7 ± 0.2 (22) |
| DPPG<br>(45°C) | Exp | 0 | 36.7 | 36.9 | 27.9 | 64.7 ± 1.0 |
|  | Sim | 0 | 37.22 ± 0.05 | 37.10 ± 0.08 | 28.61 ± 0.03 | 63.8 ± 0.2 (97) |
|  |  | 7.5 | 38.01 ± 0.04 | 37.70 ± 0.07 | 29.40 ± 0.04 | 62.7 ± 0.1 (35) |
|  |  | 15 | 38.81 ± 0.11 | 38.30 ± 0.07 | 30.21 ± 0.08 | 62.4 ± 0.3 (12) |

### Effect of STX on Bilayer Structure

We further inspected the EDPs obtained from the simulations. STX contributes with a relatively constant electron density across the inner headgroup and the hydrocarbon region of both lipids which is proportional to the STX concentration (see increase in density for the STX groups and the corresponding reduction in the density of the PG groups in Fig. 3a upon increase in STX concentration). We next split the density into the contributions of each component group, separately for the PG (Fig. S4a,b) and STX (Fig. 3b–d) lipids. The distribution of the PG groups remained largely unaffected by the presence of STX (Fig. S4a,b). Nevertheless, the STX moieties accommodated differently in response to the PG surrounding lipid and STX concentration (Fig. 3b,c). The STX headgroups (blue) aligned underneath and slightly more closely to the DMPG headgroup region (centered around dashed line), compared to those in DPPG. This is likely due to DMPG having a smaller thickness, which limits space for STX in the remainder of the bilayer. Likewise, the most prevalent location of the STX conjugated chain (magenta) was in the hydrocarbon region, below the PG headgroups. Such groups accommodated there, more tightly packed in the DMPG bilayer, exhibiting a higher density and two pronounced peaks in the distribution, as compared to the more smeared out and lower in magnitude distribution in the DPPG bilayer. The terminal methyl group densities of the STX conjugated (green) and acyl chains (red) shifted to either side of the bilayer center, displaying a large positional variance, which was dependent both on the STX concentration and the surrounding bulk PG lipid. For both lipid bilayers, it is evident that the STX conjugated chain spans both monolayers, with the terminal methyl groups occasionally extending into the opposing headgroup of PG lipids (Fig. 3d). The role of such interdigitation for the formation of entangled aggregates will be investigated in detail below.

**Figure 3.**
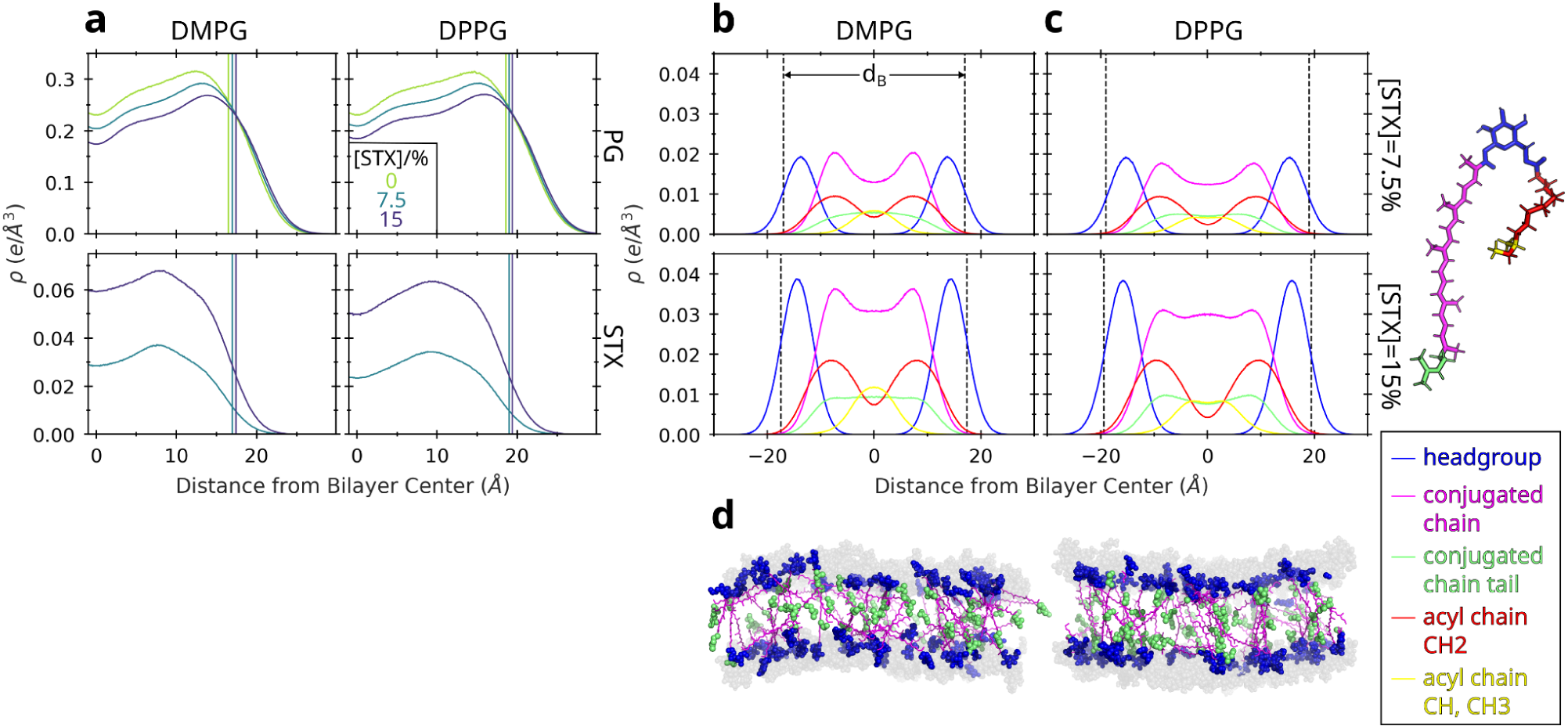
Electron density profiles of STX were recovered from MD simulations of STX–PG lipid bilayer mixtures. **(a)** Densities of PG and STX lipids (rows) for bilayers containing either DMPG or DMPG (columns) at the indicated STX concentration (color). To guide the eye, the vertical solid lines depict the density point related to the thickness d_B_ (see b–c). Note that STX electron density almost doubled upon increase in STX concentration. **(b–c)** Density profiles of STX component groups with (c) DMPG and (d) DPPG lipids at the indicated STX concentration (rows). The moieties are colored according to the legend at the right. The bilayer thickness, d_B_, was obtained from the decay in the water density at positions indicated with the vertical dashed lines (shown here to guide the eye). **(d)** Visualization of the STX molecules in the bilayer, from a typical conformation obtained from the simulations, at 15% mol STX, for DMPG (left) and DPPG (right). The PG lipid headgroups are shown in dark gray and the STX acyl-chain groups are omitted for clarity. See density profiles for the PG component groups in figure S4. Water is not shown.

### Staphyloxanthin increases the lipid packing

We next investigated the impact of STX on the lipid packing. Remarkably, the form factors revealed that lobe position shifted to lower q values as the concentration of STX increased, both in X-ray scattering experiments (Fig. 2a) and in simulations (Fig. 2b). This shift is consistent with membrane thickening. Indeed, the presence of STX increased the bilayer thickness, independently of the type of PG lipid, either DMPG or DPPG (Figs. 4a, S4c and Table 2). The observed smaller thickness for DMPG is consistent with its shorter hydrocarbon chain (14 C’s) compared with DPPG (16 C’s). For pure PG lipid bilayers, both by XDS and simulations, we obtained thickness values that are very close to previous experimental estimates (58). More importantly, when adding STX, we observed a significant increase of almost 2 Å in thickness. We also checked the lateral packing of the lipids in the MD simulations (Fig. 4b, Table 2). By measuring the area per lipid we obtained values of 0.665±0.002 nm^2^ for DMPG and 0.647±0.010 nm^2^ for DPPG. These new estimates were close to previous estimates (58). Our estimates from the simulations were lower but still very close to these experimental estimates. Reassuringly, the thickening of the bilayer, due to the presence of STX, was accompanied by a reduction in the area per lipid (Fig. 4a–b and Table 2). We also inspected the level of lipid packing locally. As seen also globally (Fig. 3), the STX molecules located underneath the PG lipids and both of them increased their local thickness as the STX concentration was increased (Fig. S5a). Interestingly, we observed that STX lipids were laterally much more tightly packed than the PG molecules (Fig. S5c). Nevertheless, the PG lipids reduced their lateral area (Fig. S5d), and moderately modified their thickness, as they got closer to the STX lipids (Fig. S5b). Thus, the observed lateral increase in lipid packing upon change in STX concentration is not uniform, but it appears to be a consequence of the local reduction in lateral area per lipid near the STX molecules. In addition, we examined the average orientation of the surrounding bulk lipids by computing the deuterium order parameter from the MD trajectories. In response to the increased lateral density and thickening, the bulk PG lipids were found to be much more orderly aligned to the membrane normal when STX was present in the membrane (Fig. 4c).

**Figure 4:**
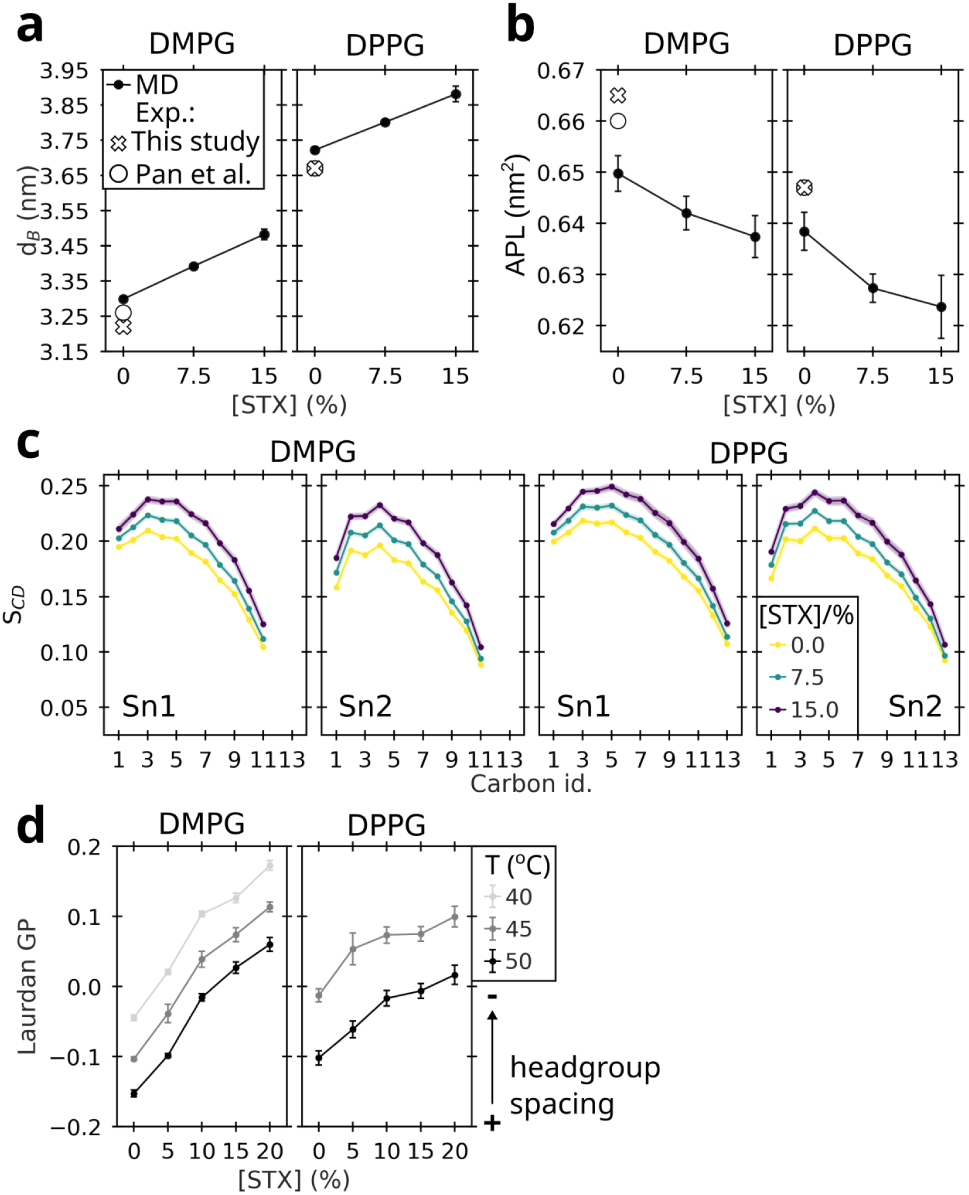
Staphyloxanthin (STX) increases lipid packing in STX–PG lipid bilayers. (a–b) Bilayer thickness, d_B_ (derived from water density) (a) and area per lipid (b) are shown as a function of the STX concentration for the indicated PG bulk lipids. Experimental values for the pure PG bilayers were obtained from X-ray scattering measurements (crosses). Previous experimental values from Pan et al. (58) are also shown. Estimates from MD simulations correspond to average ± s.e. (see values in table 2). **(c)** Deuterium order parameters, S_CD_, for both acyl chains (Sn1 and Sn2) of the indicated PG lipids were extracted from the simulations of STX-PG bilayers at the indicated STX concentration (color). **(d)** Laurdan generalized polarization is plotted as a function of the STX concentration when the bilayer was in the fluid phase at the indicated temperatures (gray shades) (average ± s.d., n=4). Note that an increase in Laurdan GP values corresponds to a decrease in headgroup spacing.

We also examined the level of headgroup spacing, through fluorescence spectroscopy. First, we reproduced our previous finding (28) that increasing STX concentration augments the Laurdan generalized polarization (GP) in DMPG lipid bilayers, indicating a reduction in head-group separation (Figs. 4d, S6a). Next, we extended this observation to DPPG, a lipid with a longer acyl chain (Figs. 4d, S6a). Note that this effect was evident at temperatures in which the bilayers were in the fluid phase, including physiologically-relevant temperatures in the case of DMPG (Fig. S6a).

Thus, our MD simulations combined with our x-ray scattering, and Laurdan experiments demonstrate that staphyloxanthin increases the lipid packing of PG lipid bilayers.

### Staphyloxanthin entangles forming trellis-like domain structures that span both leaflets

We inspected more closely the structural arrangement of staphyloxanthin. We focused on its conjugated chain. As mentioned above, the acyl chain of STX fits in, as expected, in a manner similar to the acyl chains of the bulk PG lipids (compare EDPs for acyl chains in Figs. 3 and S4. On the contrary, due to its length and rigidity, the conjugated chain displayed a distinct electron density profile (Fig. 3b–c). This chain adopted a broad range of orientations, ranging from orthogonal to-the-bilayer-surface configurations up to diagonal or even parallel to-the-bilayer-surface configurations (Fig. 5a). Note that the orientation distribution was found to depend on the length of the surrounding bulk PG lipids and the STX concentration. Accordingly, at 15 % mol concentration, STX favoured more orthogonal orientations and this effect was more pronounced for DPPG as compared to DMPG. By adopting orthogonal configurations, the conjugated chains were highly interdigitated (Figs. 5b, S7a). More generally, the extent of contact of the STX molecules, taking into account leaflet interdigitation (of the conjugated chains) and also lateral contacts, was evidenced by the formation of STX clusters (Fig. 5c). We quantified the degree of clustering by the contact area between clustered molecules, i.e. the larger this area the higher the extent of clustering (Fig. 5c). The contact area between clustered molecules was consistently dependent on the STX concentration, taking the largest values (large level of clustering) at [STX]=15 %mol (Fig. 5c). The type of PG lipid slightly influenced the contact area between STX molecules, with the thinner DMPG bilayers promoting a higher level of clustering by increased contact areas (Fig. 5c). This is consistent with the more diagonal (and even horizontal configurations) the conjugated chain of STX adopted when surrounded by this shorter type of lipid (Fig. 5a). Note that another metric to quantify the level of clustering, i.e. the size (in number of lipids) of the largest cluster, followed the same trend as the contact area (Fig. S7b). Although it should be noted this quantity is more sensitive to the transient change in inter-atomic contacts and thus less robust than the contact area.

**Figure 5:**
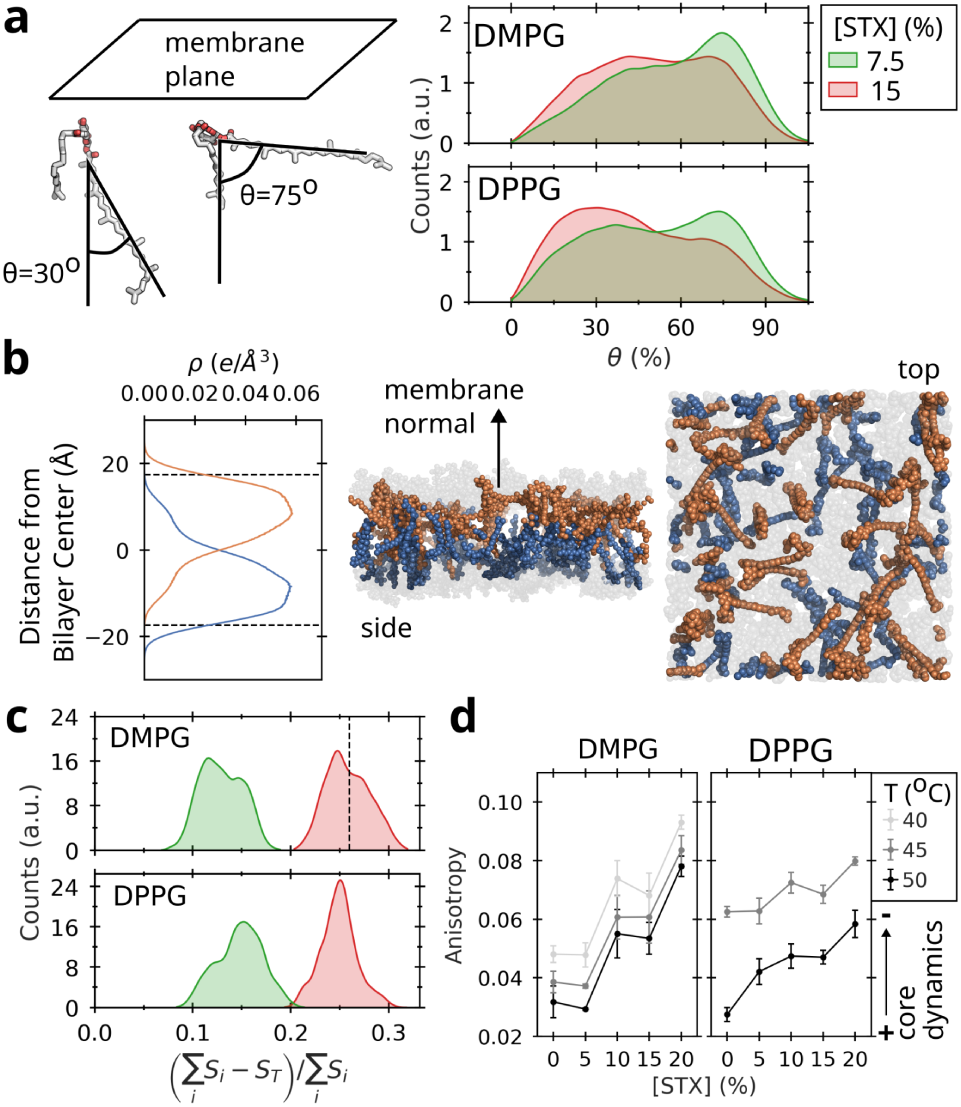
Staphyloxanthin entangles forming clusters that span both leaflets. **(a)** Orientation angle, θ, of the STX conjugated chain with respect to the vector orthogonal to membrane plane was monitored for the indicated PG bulk lipids (panels) and STX concentration (color) (distributions are shown as kernel density estimates). Examples of two orientations (θ=30° and θ=75°) are shown at the left. **(b)** (Left) electron density of STX is depicted separately here for each leaflet. The dashed lines indicate the bilayer span, according to the d_B_ thickness estimate. (Right) An example of a conformation of the STX molecules at each leaflet is also shown. To guide the eye, the PG headgroup positions (gray) and the direction of the membrane normal are shown. The density and the snapshot were taken from a simulation of the STX–DMPG bilayer at 15% mol STX concentration (see results for other concentrations and for DPPG in Fig. S7a). **(c)** Ratio of contact lipid areas, (∑_i_S_i_ - S_T_)/∑_i_S_i_, are shown, with ∑_i_S_i_ the sum of the individual surface areas of the STX molecules and S_T_ the exposed area of all STX clusters. This ratio is zero when the STX lipids are not in contact and its increase is related to the extent of formation of STX clusters. As reference, the snapshots shown in b correspond to a ratio of 0.26 (dashed line). Here, distributions are shown as kernel density estimates and colored according to the STX concentration as in a. **(d)** Anisotropy was obtained as a function of the STX concentration by DPH measurements for the indicated PG bulk lipid (panels) at temperatures in which the bilayer was in the liquid fluid phase (gray shades) (average ± s.d., n=4). Note that an increase in anisotropy corresponds to a reduction in the lipid core dynamics.

We assessed the dynamic properties of the hydrophobic core region of the membranes experimentally by performing DPH fluorescence anisotropy measurements. Here, we also first recapitulated our previous results for DMPG (28) and next assessed the behavior for DPPG. The increase in DPH anisotropy upon addition of STX indicates reduced rotational mobility of the DPH probe at the core of the lipid bilayer (Figs. 5d, 6b). This effect is more accentuated in the case of DMPG, due to the higher levels of STX network formation for this lipid. Overall, the increased restriction in the rotation of DPH is consistent with the high level of interdigitation and clustering of STX (Fig. 5a–c) and the increased ordering of the PG acyl chains (Fig. 4c) observed in the simulations.

All together, both our simulations and experiments suggest that STX forms concentration-dependent trellis-like domain entanglements that span both leaflets, impacting the structure and dynamics at the core of the lipid bilayer.

### A reduction in the gel to liquid-crystalline phase transition temperature promoted by STX is general to PG lipids of varying length

We have previously shown that the effect of STX on the phase transition temperature of DMPG lipid bilayers is only moderate (28). In contrast, by measuring CH_2_ symmetric stretching vibration, DPPG bilayers showed a more pronounced effect on the phase transition temperature due to the presence of STX. In comparison to DMPG, DPPG displayed a sharp and highly cooperative phase transition at 41.2 °C, consistent with tightly packed saturated phospholipid chains in the gel phase (Figs. S6c,d). Increasing concentrations of STX progressively shifted the transition toward lower temperatures and broadened the transition region. A downward shift in the phase transition temperature indicates a strong disruption in the acyl chain packing of the gel phase regime. The reduction in the phase transition temperature indicates that STX disrupts acyl chain packing of DPPG in the gel phase regime, reducing the stability of the gel phase and promoting an onset of phase transition at lower temperatures. Thus, STX does not only influence the structure and dynamics, but also the phase behavior of STX–PG lipid bilayers, distinctly depending on the length of the surrounding PG lipids.

## Discussion

The protective role of STX has mainly been attributed to its antioxidant activity (13). Remarkably, mechanical stabilization of the bacterial membrane by STX has also been suggested to play an important role in improving bacterial survival from host defense antimicrobial activity (26). However, an atomistic understanding of the influence of STX on the structure of lipid bilayers along with an experimental validation has been lacking. Here, we addressed this issue by a combined approach using MD simulations, X-ray scattering, fluorescence spectroscopy, and Fourier transform infrared spectroscopy..

Exploring the behavior of STX in lipid bilayers computationally by MD simulations required the parametrization of this molecule. STX was recently parameterized at an atomistic level, using the Charmm27 force-field, and at a coarse-grained level of resolution employing the MARTINI model (2.2. version) (33). Note that in this study the atomistic parameters were not validated against experimental data, but served as a reference for the coarse-grained validation. The validation was therefore indirect, based on the consistency between the all-atom and coarse-grained simulations. We provide here an alternative set of atomistic force field parameters, which is based on CGenFF. This set was optimized by a QM dihedral scan, to thereby take into account the reduced torsional mobility around the conjugated double bond connecting the triterpenoic moiety and its attachment to the glucose headgroup (Fig. S1). More importantly, the resulting parameters were validated experimentally by comparing form factors generated from the simulations with those measured by low-angle XDS. We observe an excellent agreement between the experimental and the simulated form factors (Fig. 2). This reassured that our force field MD parameters correctly reflect the molecular behavior of STX (Fig. 2). We anticipate these parameters to be highly applicable in future computational studies of STX, to improve existing all-atom models of the *S. aureus* membrane (72), which lack so far STX, and to serve as reference for refinement of coarse-grained simulations (33). Therefore, we have made them fully available for its further use and refinement at the site https://doi.org/10.17617/3.TTF8IJ.

Next, by combining simulations and experiments, the effects of STX on the structure of phosphatidylglycerol lipid bilayers were explored. We demonstrate that, in the liquid-crystalline phase, the presence of STX increases the lipid packing, as manifested by an augmentation in bilayer thickness (Figs. 2a, 4a), reduction in headgroup spacing (Figs. 4b,d), an increase in acyl chain order (Fig 4c), and a drop in mobility in the hydrophobic core (Fig. 5d). These results are in line with our previous bulk fluorescence measurements showing increased dehydration levels of the headgroup region, and reduced mobility of the hydrophobic core of DMPG bilayers in the presence of STX (28). Remarkably, we did not only recapitulate these effects for membranes containing DMPG, but we also show here that STX influence extends to longer phosphatidylglycerol lipids such as DPPG.

The triterpenoid chain is a unique structural attribute of STX. Due to its length, this chain extended into the opposite monolayer and thus induced strong levels of interdigitation (Fig. 5b), while the acyl chain was found to distribute in a manner similar to the acyl chains of the phosphatidylglycerol lipids (Fig. 3). This triterpenoid displayed a wide orientational freedom spanning configurations, that go from fully horizontal with no interdigitation to interdigitated vertical (orthogonal to the membrane plane) configurations, with a preference for the latter when the STX concentration was increased (Fig. 5a). The pronounced interdigitation and rotational freedom induced the formation of dynamic trellis-like domain structures throughout the bilayer (Fig. 5b,c). Therefore, we attribute the increase in DPH anisotropy observed in the presence of STX (Fig. 5d), which indicates a reduction in the rotational dynamics of the fluorescent probe, to not only the increase in acyl chain order of the phosphatidylglycerol groups (Fig. 4c), but also to the rotational steric hindrance induced by the presence of such STX entangled domains. This is particularly evident for DMPG bilayers, for which the DPH anisotropy is affected more drastically by the presence of STX (Fig. 5d) given by a more horizontal angle distribution of the triterpenoids (Fig. 5a) leading to a larger cluster formation (Fig. 5C), despite STX affecting the acyl chain order parameter for DMPG and DPPG at a similar level (Fig. 4c). Polar carotenoids (Xanthophylls) have been found to orient, forming an angle of about ∼40 deg with respect to the membrane normal, an orientation that is highly dictated by the positioning of their terminal hydroxyl groups at the headgroups of opposing leaflets and the connecting conjugated chain at the acyl chains region (73). In contrast, STX features only one polar terminal head group and thus the orientation of its triterpenoid is not as restricted as for Xanthophylls (Fig. 5a). We suggest that such entangled configurations of the triterpenoid chains could potentially play a role in the radical scavenging function of Staphyloxanthin.

In fluctuation experiments using giant unilamellar vesicles, *S. aureus* carotenoids have been shown to increase the bending rigidity of lipid bilayers (25). Additionally, the presence of carotenoids in model membranes increased the peptide to lipid ratio necessary to make pores (10). Moreover, an increase in lipid packing has been generally attributed to prevent the formation of pores across lipid bilayers (74, 75), as specifically demonstrated for Cardiolipin:PG mixtures (76). Thus, our results on lipid packing (Fig. 4) and clustering (Fig. 5) add a possible molecular picture of how the presence of STX increases mechanical stability of PG lipid bilayers could potentially prevent pore formation.

Carotenoid-enriched regions in *S. aureus* membranes form functional microdomains that recruit various proteins, including the penicilin-resistance factor PBP2a (30), and whose disassembly inhibits antibiotic resistance (31). Whether these functional microdomains are driven by lipid phase separation or induced through protein aggregation remains an open question. A recent combined coarse-grained MD study on membranes containing DPPG lipids, STX, and cardiolipin showed the capacity to form STX-enriched domains (33). In this study, the STX-enriched regions displayed an increased membrane thickness compared to the bulk DPPG regions. In turn, DPPG lipids increased their acyl-chain order when they were in proximity to STX molecules. Consistent with this study, in our simulations, the lipid packing was not found to be homogeneous but it varied locally. The small glucose headgroup of STX was significantly more packed, laterally, than the PG headgroups (Fig. S5c). In addition, the packing of PG molecules showed a distance dependence, with the PG molecules in the neighborhood of STX presenting a reduced area per lipid compared to bulk ones (Fig. S5d). An additional structural factor that could drive the formation of STX-enriched regions is the entanglement of the STX triterpenoid groups. In fact, the characteristic long straight triterpenoid chains and their broad angular distribution increased the probability of contact between neighboring STX molecules, leading to a clustering effect (Fig. 5b and c). Note that although the simulated spatial and temporal scale in our study was not sufficient to evidence full and reversibly the cluster formation, the local scale properties captured in our simulations—largely supported by both X-ray diffraction and fluorescence experiments—may correspond to the initial clustering events. Thus, the physicochemical properties observed here add support to the formation of STX-enriched microdomains.

*S. aureus* synthesizes varying proportions of saturated linear and branched chains (25, 77). These saturated lipids allow the microorganism to withstand oxidative stress (78, 79). Interestingly, *S. aureus* has been also suggested to sequester unsaturated acyl chains from the host (77, 80, 81). We explored the effects of the length of the acyl chains of the PG lipids. We recapitulated the gel to liquid-crystalline phase transition temperature drop in the presence of STX, observed in other model systems with STX or total *S. aureus* carotenoids (11, 33), extending it to PG lipids of varying length (Fig S6). The drop in T_m_ indicates a significant alteration of the gel phase by STX that is concomitant to increased acyl chain order in the liquid-crystalline phase. We have recently shown that the precursor 4,4’DNPA, which lacks the acyl chain and glucose moiety of STX, induces an increase in the phase transition temperature (82). The presence of the acyl chain in STX is therefore key in inducing the reduction in the phase transition temperature. Presumably, the asymmetry in length and flexibility of this side chain, in comparison to the rigid triterpenoid group, impedes the stabilization of the gel phase, while still promoting acyl-chain ordering in the fluid phase. Functionally, control of the carotenoid content has been proposed to allow *Staphylococci* to grow at low temperatures (83). Reduction in the phase transition temperature appears as a suitable mechanism by which carotenoids could help these bacteria grow in this regime. Interestingly, a larger shift in the phase transition is observed for DPPG compared to DMPG. The shift in the phase transition temperature is coupled with a widening of the liquid-crystalline/gel phase coexistence regime. This may be an indication of a possible phase coexistence that is more prevalent in the DPPG/STX mixtures. Either giant unilamellar vesicles, supported lipid bilayers or spectroscopic techniques could be used for the detection of this possible phase coexistence.

## Conclusion

By using simulations and experiments, we demonstrate that the carotenoid STX significantly alters the structure and phase behavior of phosphatidylglycerol lipid bilayers of varying lengths—specifically DMPG and DPPG. In the liquid-crystalline phase, STX enhances lipid packing and increases acyl-chain order, while lowering the transition temperature to the gel phase. By forming entangled trellis-like domain structures that span both leaflets, the conjugated triterpenoid chain of this lipid appears to be crucial for these structural changes. Such interleaflet entanglements may constitute a critical structural prerequisite for STX’s radical scavenging function in *S. aureus* membranes. Furthermore, our findings are consistent with the existence of structurally-distinct STX-enriched microdomains. It will be highly interesting to explore how these STX-induced structural changes correlate with its antioxidant function and potentially with the prevention of membrane pore formation, as survival strategies of *S aureus* to cope with stressful environmental changes.

## Supporting Information Description

Supporting information includes Figures S1–S7.

## Author contributions

D.R.F.B., J.M.D., J.D.O., J.E.C., and C.A.-S. performed the MD simulations. D.R.F.B. and G.P.M. carried out the QM dihedral scan. A.B. performed the Laurdan and DPH experiments. L.H., J.J., K.J., L.K., and S.T.-N. carried out the X-ray scattering experiments. E.S., G.-D.L., and C.C. generated the STX samples. M.M.-M. conducted the FTIR experiments. C.L. and C.A.-S. conceived and administered the project. All authors analyzed and discussed the results. D.R.F.B., A.B., L.H., J.J., G.-D.L., M.M.-M., S.T.-N., C.L., and C.A.-S. wrote the manuscript.

## Acknowledgements

We thank James C. Gumbart for sharing scripts for the calculation of Atomic electron density profiles. We are grateful for financial support by the Universidad de los Andes (C.L.), the Ministerio de Ciencia Tecnologia e Innovación de Colombia (Grant 1204-937-101846) (C.L.), and the University of Antioquia (Grant 2024–66871) (M.M.-M.). This work is also based upon research conducted at the Center for High Energy X-ray Sciences (CHEXS at CHESS), which is supported by the National Science Foundation (award DMR-1829070), and the Macromolecular Diffraction at CHESS (MacCHESS) facility, which is supported by award 1-P30-GM124166-01A1 from the National Institute of General Medical Sciences, National Institutes of Health, and by New York State’s Empire State Development Corporation. Additional support is from the National Institute of Allergy and Infectious Diseases (award 1R01AI172861-01A1) (S.T-N.) and NASA Pennsylvania Space Grant Consortium (L.K.). Computations were performed on the HPC system at the Max Planck Computing and Data Facility.

## Data availability

Relevant data are within the manuscript and its Supporting Material. Molecular dynamics simulation data and newly-generated force-field parameters of STX have been deposited on EDMOND at https://doi.org/10.17617/3.TTF8IJ.

## References

1. Laux, C., A. Peschel, and B. Krismer. 2019. Staphylococcus aureus Colonization of the Human Nose and Interaction with Other Microbiome Members. Microbiol. Spectr. 7:10.1128/microbiolspec.gpp3-0029-2018, doi: 10.1128/microbiolspec.gpp3-0029-2018.

2. Piewngam, P., and M. Otto. 2024. Staphylococcus aureus colonisation and strategies for decolonisation. Lancet Microbe 5:e606–e618, doi: 10.1016/S2666-5247(24)00040-5.

3. van Hal, S.J., S.O. Jensen, V.L. Vaska, B.A. Espedido, D.L. Paterson, and I.B. Gosbell. 2012. Predictors of Mortality in Staphylococcus aureus Bacteremia. Clin. Microbiol. Rev. 25:362–386, doi: 10.1128/cmr.05022-11.

4. Diekema, D.J., M.A. Pfaller, D. Shortridge, M. Zervos, and R.N. Jones. 2019. Twenty-Year Trends in Antimicrobial Susceptibilities Among Staphylococcus aureus From the SENTRY Antimicrobial Surveillance Program. Open Forum Infect. Dis. 6:S47–S53, doi: 10.1093/ofid/ofy270.

5. Bai, A.D., C.K. Lo, A.S. Komorowski, M. Suresh, K. Guo, A. Garg, P. Tandon, J. Senecal, O.D. Corpo, I. Stefanova, C. Fogarty, G. Butler-Laporte, E.G. McDonald, M.P. Cheng, A.M. Morris, M. Loeb, and T.C. Lee. 2022. Staphylococcus aureus bacteremia mortality across country income groups: A secondary analysis of a systematic review. Int. J. Infect. Dis. 122:405–411, doi: 10.1016/j.ijid.2022.06.026.

6. Holland, T.L., C. Arnold, and V.G. Fowler Jr. 2014. Clinical Management of Staphylococcus aureus Bacteremia: A Review. JAMA 312:1330–1341, doi: 10.1001/jama.2014.9743.

7. Bellis, K.L., O.M. Dissanayake, E.M. Harrison, and D. Aggarwal. 2025. Community methicillin-resistant Staphylococcus aureus outbreaks in areas of low prevalence. Clin. Microbiol. Infect. 31:182–189, doi: 10.1016/j.cmi.2024.06.006.

8. Foster, T.J. 2017. Antibiotic resistance in Staphylococcus aureus. Current status and future prospects. FEMS Microbiol. Rev. 41:430–449, doi: 10.1093/femsre/fux007.

9. Hammond, R.K., and D.C. White. 1970. Carotenoid Formation by Staphylococcus aureus. J. Bacteriol. 103:191–198, doi: 10.1128/jb.103.1.191-198.1970.

10. López, G.-D., E. Suesca, G. Álvarez-Rivera, A.E. Rosato, E. Ibáñez, A. Cifuentes, C. Leidy, and C. Carazzone. 2021. Carotenogenesis of *Staphylococcus aureus*: New insights and impact on membrane biophysical properties. Biochim. Biophys. Acta BBA - Mol. Cell Biol. Lipids 1866:158941, doi: 10.1016/j.bbalip.2021.158941.

11. Marshall, J.H., and G.J. Wilmoth. 1981. Pigments of Staphylococcus aureus, a series of triterpenoid carotenoids. J. Bacteriol. 147:900–913, doi: 10.1128/jb.147.3.900-913.1981.

12. Pelz, A., K.-P. Wieland, K. Putzbach, P. Hentschel, K. Albert, and F. Götz. 2005. Structure and Biosynthesis of Staphyloxanthin from Staphylococcus aureus*. J. Biol. Chem. 280:32493–32498, doi: 10.1074/jbc.M505070200.

13. Clauditz, A., A. Resch, K.-P. Wieland, A. Peschel, and F. Götz. 2006. Staphyloxanthin Plays a Role in the Fitness of Staphylococcus aureus and Its Ability To Cope with Oxidative Stress. Infect. Immun. 74:4950–4953, doi: 10.1128/iai.00204-06.

14. Zamudio-Chávez, L., E. Suesca, G.-D. López, C. Carazzone, M. Manrique-Moreno, and C. Leidy. 2023. Staphylococcus aureus Modulates Carotenoid and Phospholipid Content in Response to Oxygen-Restricted Growth Conditions, Triggering Changes in Membrane Biophysical Properties. Int. J. Mol. Sci. 24:14906, doi: 10.3390/ijms241914906.

15. Pannu, M.K., D.A. Hudman, N.J. Sargentini, and V.K. Singh. 2019. Role of SigB and Staphyloxanthin in Radiation Survival of Staphylococcus aureus. Curr. Microbiol. 76:70–77, doi: 10.1007/s00284-018-1586-x.

16. Bae, E., D. Kim, M. Kim, A. Kang, J. Shin, Y. Kim, and M. Shin. 2025. Master transcriptional regulator SaeS in Staphylococcus aureus contributes to staphyloxanthin biosynthesis to promote survival during invasive infection. Virulence 16:2580159, doi: 10.1080/21505594.2025.2580159.

17. Nicholas, R.O., T. Li, D. McDevitt, A. Marra, S. Sucoloski, P.L. Demarsh, and D.R. Gentry. 1999. Isolation and Characterization of a sigBDeletion Mutant of Staphylococcus aureus. Infect. Immun. 67:3667–3669, doi: 10.1128/iai.67.7.3667-3669.1999.

18. Pandey, S., G.S. Sahukhal, and M.O. Elasri. 2019. The msaABCR Operon Regulates the Response to Oxidative Stress in Staphylococcus aureus. J. Bacteriol. 201:10.1128/jb.00417-19, doi: 10.1128/jb.00417-19.

19. Austin, C.M., S. Garabaglu, C.N. Krute, M.J. Ridder, N.A. Seawell, M.A. Markiewicz, J.M. Boyd, and J.L. Bose. 2019. Contribution of YjbIH to Virulence Factor Expression and Host Colonization in Staphylococcus aureus. Infect. Immun. 87:10.1128/iai.00155-19, doi: 10.1128/iai.00155-19.

20. Hall, J.W., J. Yang, H. Guo, and Y. Ji. 2017. The Staphylococcus aureus AirSR Two-Component System Mediates Reactive Oxygen Species Resistance via Transcriptional Regulation of Staphyloxanthin Production. Infect. Immun. 85:10.1128/iai.00838-16, doi: 10.1128/iai.00838-16.

21. Yu, J., L. Shen, J. Yang, J. Shi, Y. Huang, Y. Shang, and F. Yu. 2025. Staphyloxanthin-enriched extracts promote biofilm formation and oxidative stress resistance in Staphylococcus aureus. Microbiol. Spectr. 13:e00996–25, doi: 10.1128/spectrum.00996-25.

22. Nosair, A.M., A.A. Abdelaziz, A.M. Abo-Kamar, L.A. Al-Madboly, and M.H. Farghali. 2025. Deciphering the efficacy of staphyloxanthin-encapsulated niosomal nanovesicles to attenuate biofilm formation, quorum sensing, and meropenem persistence in Acinetobacter baumannii. BMC Microbiol. 25:791, doi: 10.1186/s12866-025-04507-1.

23. Song, Y., C.-I. Liu, F.-Y. Lin, J.H. No, M. Hensler, Y.-L. Liu, W.-Y. Jeng, J. Low, G.Y. Liu, V. Nizet, A.H.-J. Wang, and E. Oldfield. 2009. Inhibition of Staphyloxanthin Virulence Factor Biosynthesis in Staphylococcus aureus: In Vitro, in Vivo, and Crystallographic Results. J. Med. Chem. 52:3869–3880, doi: 10.1021/jm9001764.

24. Fuertes-Chaves, C., J.E. Gonzalez, E. Suesca, P. Guzmán-Sastoque, C. Muñoz-Camargo, M. Manrique-Moreno, C. Carazzone, and C. Leidy. 2026. Exposure to the antimicrobial peptides LL-37 and ATRA-1 induces a lipidome response in Staphylococcus aureus that alters membrane biophysical properties. bioRxiv.

25. Perez-Lopez, M.I., R. Mendez-Reina, S. Trier, C. Herrfurth, I. Feussner, A. Bernal, M. Forero-Shelton, and C. Leidy. 2019. Variations in carotenoid content and acyl chain composition in exponential, stationary and biofilm states of *Staphylococcus aureus*, and their influence on membrane biophysical properties. Biochim. Biophys. Acta BBA - Biomembr. 1861:978–987, doi: 10.1016/j.bbamem.2019.02.001.

26. Mishra, N.N., G.Y. Liu, M.R. Yeaman, C.C. Nast, R.A. Proctor, J. McKinnell, and A.S. Bayer. 2011. Carotenoid-Related Alteration of Cell Membrane Fluidity Impacts Staphylococcus aureus Susceptibility to Host Defense Peptides. Antimicrob. Agents Chemother. 55:526–531, doi: 10.1128/aac.00680-10.

27. Manrique-Moreno, M., M. Jemioła-Rzemińska, J. Múnera-Jaramillo, G.-D. López, E. Suesca, C. Leidy, and K. Strzałka. 2022. Staphylococcus aureus Carotenoids Modulate the Thermotropic Phase Behavior of Model Systems That Mimic Its Membrane Composition. Membranes 12:945, doi: 10.3390/membranes12100945.

28. Múnera-Jaramillo, J., G.-D. López, E. Suesca, C. Carazzone, C. Leidy, and M. Manrique-Moreno. 2024. The role of staphyloxanthin in the regulation of membrane biophysical properties in *Staphylococcus aureus*. Biochim. Biophys. Acta BBA - Biomembr. 1866:184288, doi: 10.1016/j.bbamem.2024.184288.

29. White, D.C., and F.E. Frerman. 1967. Extraction, Characterization, and Cellular Localization of the Lipids of Staphylococcus aureus. J. Bacteriol. 94:1854–1867, doi: 10.1128/jb.94.6.1854-1867.1967.

30. Adolf, L.A., A. Müller-Jochim, L. Kricks, J.-S. Puls, D. Lopez, F. Grein, and S. Heilbronner. 2023. Functional membrane microdomains and the hydroxamate siderophore transporter ATPase FhuC govern Isd-dependent heme acquisition in Staphylococcus aureus. eLife 12:e85304, doi: 10.7554/eLife.85304.

31. García-Fernández, E., G. Koch, R.M. Wagner, A. Fekete, S.T. Stengel, J. Schneider, B. Mielich-Süss, S. Geibel, S.M. Markert, C. Stigloher, and D. Lopez. 2017. Membrane Microdomain Disassembly Inhibits MRSA Antibiotic Resistance. Cell 171:1354–1367.e20, doi: 10.1016/j.cell.2017.10.012.

32. Ukleja, M., L. Kricks, G. Torrens, I. Peschiera, I. Rodrigues-Lopes, M. Krupka, J. García-Fernández, R. Melero, R. del Campo, A. Eulalio, A. Mateus, M. López-Bravo, A.I. Rico, F. Cava, and D. Lopez. 2024. Flotillin-mediated stabilization of unfolded proteins in bacterial membrane microdomains. Nat. Commun. 15:5583, doi: 10.1038/s41467-024-49951-1.

33. Gupta, S., and T. Mandal. 2022. Simulation study of domain formation in a model bacterial membrane. Phys. Chem. Chem. Phys. 24:18133–18143, doi: 10.1039/D2CP01873J.

34. Vanommeslaeghe, K., E. Hatcher, C. Acharya, S. Kundu, S. Zhong, J. Shim, E. Darian, O. Guvench, P. Lopes, I. Vorobyov, and A.D. Mackerell Jr. 2010. CHARMM general force field: A force field for drug-like molecules compatible with the CHARMM all-atom additive biological force fields. J. Comput. Chem. 31:671–690, doi: 10.1002/jcc.21367.

35. Jo, S., T. Kim, V.G. Iyer, and W. Im. 2008. CHARMM-GUI: A Web-Based Graphical User Interface for CHARMM. J. Comput. Chem. 29:1859–1865, doi: 10.1002/jcc.20945.

36. Frisch, M.J., G.W. Trucks, H.B. Schlegel, G.E. Scuseria, M.A. Robb, J.R. Cheeseman, G. Scalmani, V. Barone, G.A. Petersson, H. Nakatsuji, X. Li, M. Caricato, A.V. Marenich, J. Bloino, B.G. Janesko, R. Gomperts, B. Mennucci, H.P. Hratchian, J.V. Ortiz, A.F. Izmaylov, J.L. Sonnenberg, D. Williams-Young, F. Ding, F. Lipparini, F. Egidi, J. Goings, B. Peng, A. Petrone, T. Henderson, D. Ranasinghe, V.G. Zakrzewski, J. Gao, N. Rega, G. Zheng, W. Liang, M. Hada, M. Ehara, K. Toyota, R. Fukuda, J. Hasegawa, M. Ishida, T. Nakajima, Y. Honda, O. Kitao, H. Nakai, T. Vreven, K. Throssell, J.A. Montgomery Jr., J.E. Peralta, F. Ogliaro, M.J. Bearpark, J.J. Heyd, E.N. Brothers, K.N. Kudin, V.N. Staroverov, T.A. Keith, R. Kobayashi, J. Normand, K. Raghavachari, A.P. Rendell, J.C. Burant, S.S. Iyengar, J. Tomasi, M. Cossi, J.M. Millam, M. Klene, C. Adamo, R. Cammi, J.W. Ochterski, R.L. Martin, K. Morokuma, O. Farkas, J.B. Foresman, and D.J. Fox. 2016. Gaussiañ16 Revision C.01.

37. Requena, A., J.P. Cerón-Carrasco, A. Bastida, J. Zúñiga, and B. Miguel. 2008. A Density Functional Theory Study of the Structure and Vibrational Spectra of β-Carotene, Capsanthin, and Capsorubin. J. Phys. Chem. A 112:4815–4825, doi: 10.1021/jp710304u.

38. Klauda, J.B., V. Monje, T. Kim, and W. Im. 2012. Improving the CHARMM Force Field for Polyunsaturated Fatty Acid Chains. J. Phys. Chem. B 116:9424–9431, doi: 10.1021/jp304056p.

39. Klauda, J.B., R.M. Venable, J.A. Freites, J.W. O’Connor, D.J. Tobias, C. Mondragon-Ramirez, I. Vorobyov, A.D.Jr. MacKerell, and R.W. Pastor. 2010. Update of the CHARMM All-Atom Additive Force Field for Lipids: Validation on Six Lipid Types. J. Phys. Chem. B 114:7830–7843, doi: 10.1021/jp101759q.

40. MacKerell, A.D.Jr., D. Bashford, M. Bellott, R.L.Jr. Dunbrack, J.D. Evanseck, M.J. Field, S. Fischer, J. Gao, H. Guo, S. Ha, D. Joseph-McCarthy, L. Kuchnir, K. Kuczera, F.T.K. Lau, C. Mattos, S. Michnick, T. Ngo, D.T. Nguyen, B. Prodhom, W.E. Reiher, B. Roux, M. Schlenkrich, J.C. Smith, R. Stote, J. Straub, M. Watanabe, J. Wiórkiewicz-Kuczera, D. Yin, and M. Karplus. 1998. All-Atom Empirical Potential for Molecular Modeling and Dynamics Studies of Proteins. J. Phys. Chem. B 102:3586–3616, doi: 10.1021/jp973084f.

41. Jorgensen, W.L., J. Chandrasekhar, J.D. Madura, R.W. Impey, and M.L. Klein. 1983. Comparison of simple potential functions for simulating liquid water. J. Chem. Phys. 79:926–935, doi: 10.1063/1.445869.

42. Beglov, D., and B. Roux. 1994. Finite representation of an infinite bulk system: Solvent boundary potential for computer simulations. J. Chem. Phys. 100:9050–9063, doi: 10.1063/1.466711.

43. Won, Y. 2012. Force Field for Monovalent, Divalent, and Trivalent Cations Developed under the Solvent Boundary Potential. J. Phys. Chem. A 116:11763–11767, doi: 10.1021/jp309150r.

44. Nosé, S. 1984. A Unified Formulation of the Constant Temperature Molecular Dynamics Methods. J. Chem. Phys. 81:511–519, doi: 10.1063/1.447334.

45. Parrinello, M., and A. Rahman. 1981. Polymorphic transitions in single crystals: A new molecular dynamics method. J. Appl. Phys. 52:7182–7190, doi: 10.1063/1.328693.

46. Darden, T., D. York, and L. Pedersen. 1993. Particle mesh Ewald: An N⋅log(N) method for Ewald sums in large systems. J. Chem. Phys. 98:10089–10092, doi: 10.1063/1.464397.

47. Essmann, U., L. Perera, M.L. Berkowitz, T. Darden, H. Lee, and L.G. Pedersen. 1995. A smooth particle mesh Ewald method. J. Chem. Phys. 103:8577–8593, doi: 10.1063/1.470117.

48. Páll, S., and B. Hess. 2013. A flexible algorithm for calculating pair interactions on SIMD architectures. Comput. Phys. Commun. 184:2641–2650, doi: 10.1016/j.cpc.2013.06.003.

49. Hess, B., H. Bekker, H.J.C. Berendsen, and J.G.E.M. Fraaije. 1997. LINCS: A Linear Constraint Solver for Molecular Simulations. J. Comput. Chem. 18:1463–1472, doi: 10.1002/(SICI)1096-987X(199709)18:12<1463::AID-JCC4>3.0.CO;2-H.

50. Miyamoto, S., and P.A. Kollman. 1992. Settle: An analytical version of the SHAKE and RATTLE algorithm for rigid water models. J. Comput. Chem. 13:952–962, doi: 10.1002/jcc.540130805.

51. Abraham, M.J., T. Murtola, R. Schulz, S. Páll, J.C. Smith, B. Hess, and E. Lindahl. 2015. GROMACS: High performance molecular simulations through multi-level parallelism from laptops to supercomputers. SoftwareX 1–2:19–25, doi: 10.1016/j.softx.2015.06.001.

52. Michaud-Agrawal, N., E.J. Denning, T.B. Woolf, and O. Beckstein. 2011. MDAnalysis: A toolkit for the analysis of molecular dynamics simulations. J. Comput. Chem. 32:2319–2327, doi: 10.1002/jcc.21787.

53. Brandt, E.G., A.R. Braun, J.N. Sachs, J.F. Nagle, and O. Edholm. 2011. Interpretation of Fluctuation Spectra in Lipid Bilayer Simulations. Biophys. J. 100:2104–2111, doi: 10.1016/j.bpj.2011.03.010.

54. Braun, A.R., E.G. Brandt, O. Edholm, J.F. Nagle, and J.N. Sachs. 2011. Determination of Electron Density Profiles and Area from Simulations of Undulating Membranes. Biophys. J. 100:2112–2120, doi: 10.1016/j.bpj.2011.03.009.

55. Kučerka, N., J.F. Nagle, J.N. Sachs, S.E. Feller, J. Pencer, A. Jackson, and J. Katsaras. 2008. Lipid Bilayer Structure Determined by the Simultaneous Analysis of Neutron and X-Ray Scattering Data. Biophys. J. 95:2356–2367, doi: 10.1529/biophysj.108.132662.

56. Kučerka, N., M.-P. Nieh, and J. Katsaras. 2011. Fluid phase lipid areas and bilayer thicknesses of commonly used phosphatidylcholines as a function of temperature. Biochim. Biophys. Acta BBA - Biomembr. 1808:2761–2771, doi: 10.1016/j.bbamem.2011.07.022.

57. Pan, J., F.A. Heberle, S. Tristram-Nagle, M. Szymanski, M. Koepfinger, J. Katsaras, and N. Kučerka. 2012. Molecular structures of fluid phase phosphatidylglycerol bilayers as determined by small angle neutron and X-ray scattering. Biochim. Biophys. Acta BBA - Biomembr. 1818:2135–2148, doi: 10.1016/j.bbamem.2012.05.007.

58. Pan, J., D. Marquardt, F.A. Heberle, N. Kučerka, and J. Katsaras. 2014. Revisiting the bilayer structures of fluid phase phosphatidylglycerol lipids: Accounting for exchangeable hydrogens. Biochim. Biophys. Acta BBA - Biomembr. 1838:2966–2969, doi: 10.1016/j.bbamem.2014.08.009.

59. Petrache, H.I., S.E. Feller, and J.F. Nagle. 1997. Determination of component volumes of lipid bilayers from simulations. Biophys. J. 72:2237–2242, doi: 10.1016/S0006-3495(97)78867-2.

60. Kučerka, N., J. Katsaras, and J.F. Nagle. 2010. Comparing Membrane Simulations to Scattering Experiments: Introducing the SIMtoEXP Software. J. Membr. Biol. 235:43–50, doi: 10.1007/s00232-010-9254-5.

61. Gapsys, V., B.L. de Groot, and R. Briones. 2013. Computational analysis of local membrane properties. J. Comput. Aided Mol. Des. 27:845–858, doi: 10.1007/s10822-013-9684-0.

62. Abraham, M., A. Alekseenko, B. Andrews, P. Bauer, C. Bergh, H. Bird, E. Briand, A. Brown, Y. Chen, M. Doijade, G. Fiorin, S. Fleischmann, S. Gorelov, G. Gouaillardet, A. Gray, F. Jalalypour, P. Johansson, C. Kutzner, G. Lazarski, J. Lemkul, M. Lundborg, J. Maia, P.T. Merz, V. Miletić, D. Morozov, L. Müllender, S. Páll, A. Pasquadibisceglie, M. Pellegrino, N. Piasentin, D. Rapetti, M.U. Sadiq, H. Santuz, M. Shirts, T. Shugaeva, A. Shvetsov, B. Soproni, P. Turner, A. Villa, Y. Zhang, B. Hess, and E. Lindahl. 2026. GROMACS 2026.2 Manual. doi: 10.5281/zenodo.20037907.

63. Egberts, E., and H.J.C. Berendsen. 1988. Molecular dynamics simulation of a smectic liquid crystal with atomic detail. J. Chem. Phys. 89:3718–3732, doi: 10.1063/1.454893.

64. Landrum, G., P. Tosco, B. Kelley, R. Rodriguez, D. Cosgrove, R. Vianello, sriniker, P. Gedeck, G. Jones, D. Nealschneider, E. Kawashima, NadineSchneider, tadhurst-cdd, A. Dalke, M. Swain, B. Cole, S. Turk, A. Savelev, N. Maeder, Y. Pechersky, A. Vaucher, M. Wójcikowski, R. Walker, H. Faara, I. Take, V.F. Scalfani, D. Probst, K. Ujihara, J. Monat, and J. Lehtivarjo. 2026. rdkit/rdkit: 2026_03_3 (Q1 2026) Release. Zenodo.

65. Humphrey, W., A. Dalke, and K. Schulten. 1996. VMD: visual molecular dynamics. J. Mol. Graph. Model. 14:33–8, 27–8, doi: 10.1016/0263-7855(96)00018-5.

66. Schrödinger, LLC. 2015. The PyMOL Molecular Graphics System, Version 1.8.

67. Tristram-Nagle, S.A. 2007. Preparation of Oriented, Fully Hydrated Lipid Samples for Structure Determination Using X-Ray Scattering. In Methods in Membrane Lipids. Dopico AM, editor. Humana Press: Totowa, NJ, pp. 63–75.

68. Mitra, S., M. Coopershlyak, Y. Li, B. Chandersekhar, R. Koenig, M.-T. Chen, B. Evans, F. Heinrich, B. Deslouches, and S. Tristram-Nagle. 2023. Novel Helical Trp- and Arg-Rich Antimicrobial Peptides Locate Near Membrane Surfaces and Rigidify Lipid Model Membranes. Adv. NanoBiomed Res. 3:2300013, doi: 10.1002/anbr.202300013.

69. Lyatskaya, Y., Y. Liu, S. Tristram-Nagle, J. Katsaras, and J.F. Nagle. 2000. Method for obtaining structure and interactions from oriented lipid bilayers. *Phys*. Rev. E 63:011907, doi: 10.1103/PhysRevE.63.011907.

70. Helfrich, W. 1978. Steric Interaction of Fluid Membranes in Multilayer Systems. Z. Naturforschung - Sect. J. Phys. Sci. 33:305–315, doi: 10.1515/zna-1978-0308.

71. Gennes, P.G. de, and J. Prost. 1995. The Physics of Liquid Crystals. Second Edition, Second Edition. Oxford University Press: Oxford, New York.

72. Joodaki, F., L.M. Martin, and M.L. Greenfield. 2022. Generation and Computational Characterization of a Complex Staphylococcus aureus Lipid Bilayer. Langmuir 38:9481–9499, doi: 10.1021/acs.langmuir.2c00483.

73. Grudzinski, W., L. Nierzwicki, R. Welc, E. Reszczynska, R. Luchowski, J. Czub, and W.I. Gruszecki. 2017. Localization and Orientation of Xanthophylls in a Lipid Bilayer. Sci. Rep. 7:9619, doi: 10.1038/s41598-017-10183-7.

74. Ting, C.L., N. Awasthi, M. Müller, and J.S. Hub. 2018. Metastable Prepores in Tension-Free Lipid Bilayers. Phys. Rev. Lett. 120:128103, doi: 10.1103/PhysRevLett.120.128103.

75. Hub, J.S., and N. Awasthi. 2017. Probing a Continuous Polar Defect: A Reaction Coordinate for Pore Formation in Lipid Membranes. J. Chem. Theory Comput. 13:2352–2366, doi: 10.1021/acs.jctc.7b00106.

76. Rocha-Roa, C., J.D. Orjuela, C. Leidy, P. Cossio, and C. Aponte-Santamaría. 2021. Cardiolipin prevents pore formation in phosphatidylglycerol bacterial membrane models. FEBS Lett. 595:2701–2714, doi: 10.1002/1873-3468.14206.

77. Frank, M.W., J. Yao, J.L. Batte, J.M. Gullett, C. Subramanian, J.W. Rosch, and C.O. Rock. 2020. Host Fatty Acid Utilization by Staphylococcus aureus at the Infection Site. mBio 11:10.1128/mbio.00920-20, doi: 10.1128/mbio.00920-20.

78. Sen, S., S. Sirobhushanam, S.R. Johnson, Y. Song, R. Tefft, C. Gatto, and B.J. Wilkinson. 2016. Growth-Environment Dependent Modulation of Staphylococcus aureus Branched-Chain to Straight-Chain Fatty Acid Ratio and Incorporation of Unsaturated Fatty Acids. PLOS ONE 11:e0165300, doi: 10.1371/journal.pone.0165300.

79. Singh, V.K., D.S. Hattangady, E.S. Giotis, A.K. Singh, N.R. Chamberlain, M.K. Stuart, and B.J. Wilkinson. 2008. Insertional Inactivation of Branched-Chain α-Keto Acid Dehydrogenase in Staphylococcus aureus Leads to Decreased Branched-Chain Membrane Fatty Acid Content and Increased Susceptibility to Certain Stresses. Appl. Environ. Microbiol. 74:5882–5890, doi: 10.1128/AEM.00882-08.

80. Hines, K.M., G. Alvarado, X. Chen, C. Gatto, A. Pokorny, F. Alonzo, B.J. Wilkinson, and L. Xu. 2020. Lipidomic and Ultrastructural Characterization of the Cell Envelope of Staphylococcus aureus Grown in the Presence of Human Serum. mSphere 5:10.1128/msphere.00339-20, doi: 10.1128/msphere.00339-20.

81. Raskovic, D., G. Alvarado, K.M. Hines, L. Xu, C. Gatto, B.J. Wilkinson, and A. Pokorny. 2025. Growth of *Staphylococcus aureus* in the presence of oleic acid shifts the glycolipid fatty acid profile and increases resistance to antimicrobial peptides. Biochim. Biophys. Acta BBA - Biomembr. 1867:184395, doi: 10.1016/j.bbamem.2024.184395.

82. Múnera-Jaramillo, J., G.-D. López, E. Suesca, E. Ibáñez, A. Cifuentes, C. Carazzone, C. Leidy, and M. Manrique-Moreno. 2026. Membrane structural properties in Staphylococcus aureus are tuned by the carotenoid 4,4′-diaponeurosporenoic acid. bioRxiv.

83. Seel, W., D. Baust, D. Sons, M. Albers, L. Etzbach, J. Fuss, and A. Lipski. 2020. Carotenoids are used as regulators for membrane fluidity by Staphylococcus xylosus. Sci. Rep. 10:330, doi: 10.1038/s41598-019-57006-5.

