## Supporting information for "The *Staphylococcus aureus* carotenoid staphyloxanthin modifies the structure of phosphatidylglycerol lipid bilayers"

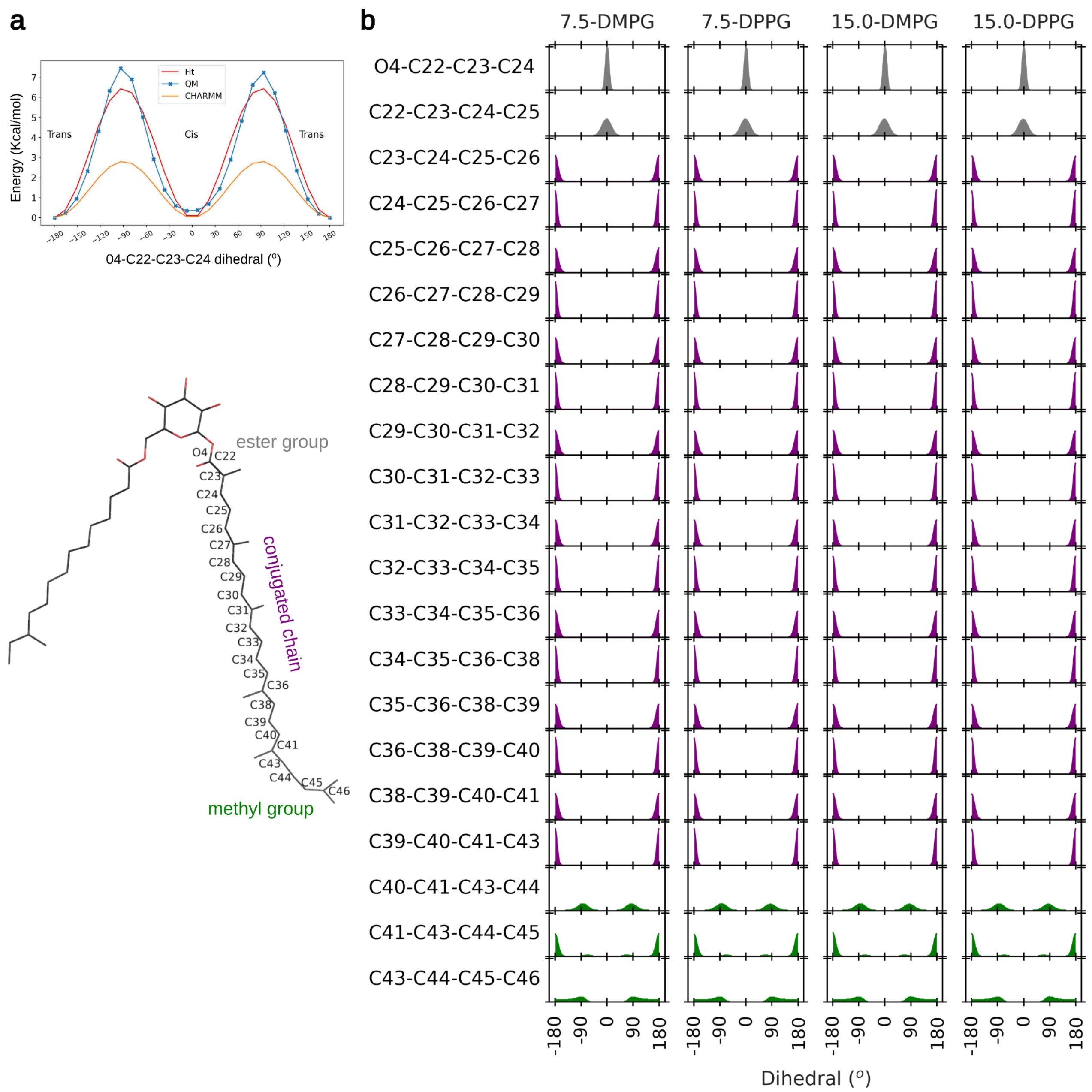


**Figure S1.** Parametrization staphyloxanthin. The CGENFF scheme was utilized for most parameters. **a)** Quantum mechanics (QM) dihedral optimization of the dihedral O4–C22–C23–C24 was carried out. Accordingly, the dihedral parameters around the C22-C23 bond were increased to fit the QM dihedral curve and thereby preserve these torsions at the right cis or trans configuration. **b)** Dihedrals along the conjugated chain (purple), the connecting ester group (gray) and the terminal methyl group (green) (rows) recovered from the equilibrium MD simulation at the indicated STX concentration (7.5 or 15 %mol) and bulk PG lipid (columns). Atoms considered in each dihedral are indicated at the left snapshot. The dihedrals confirmed that the *trans* configuration (± 180°) was maintained, highlighting the rigid nature of the conjugated chain.

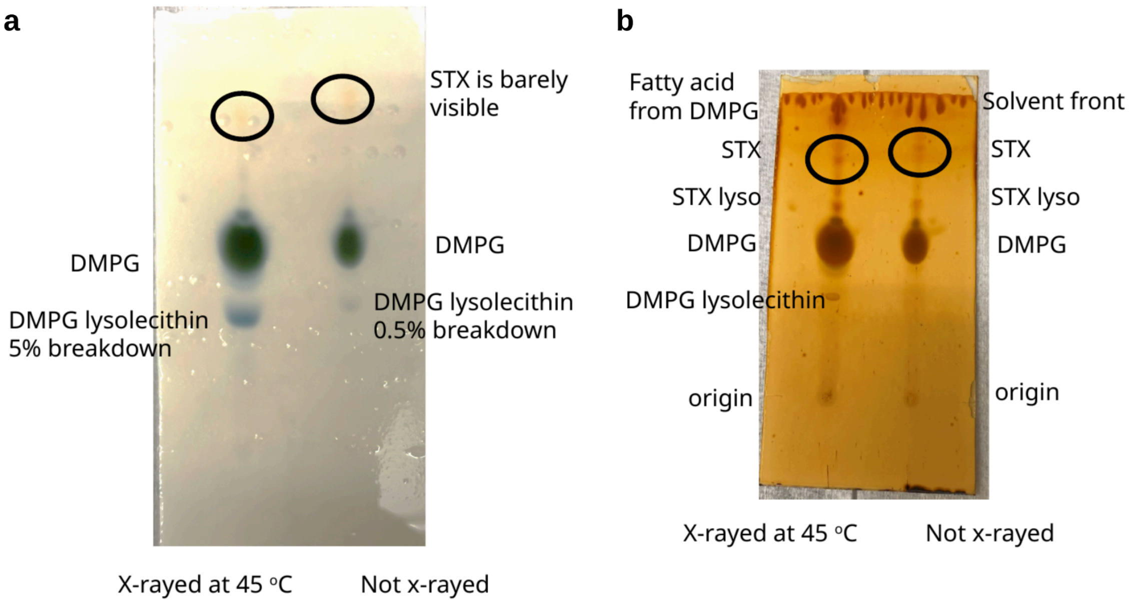


**Figure S2.** DMPG/10% STX TLC, stained with Molybdic acid which binds to phosphate **(a)** and with iodine vapor after molybdic acid is evaporated **(b)**. Percent breakdown is estimated visually from a standard curve of lysolecithin in DPPC.

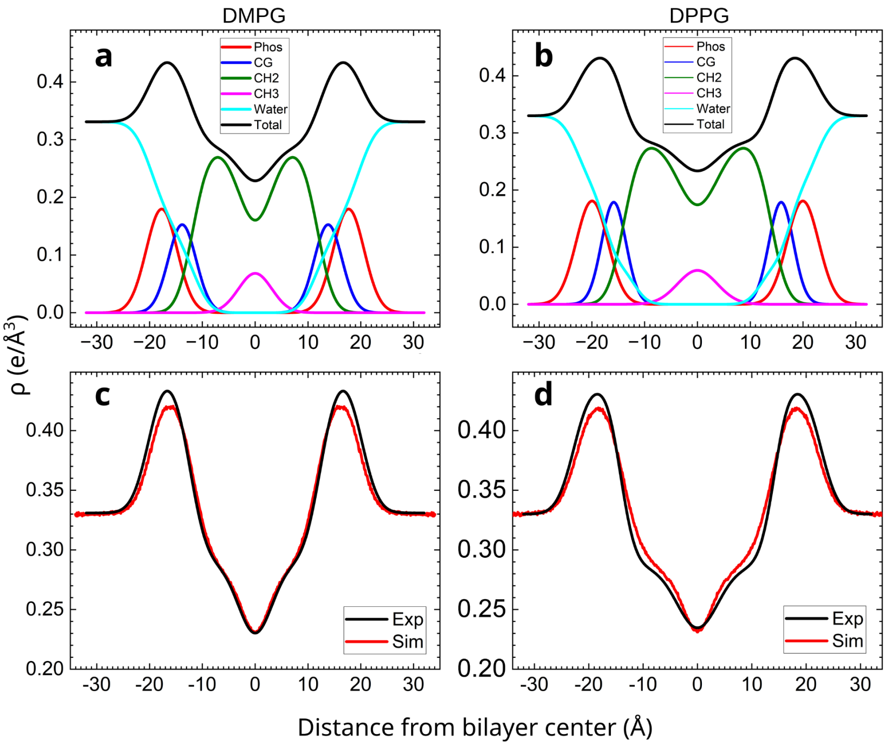


**Figure S3. Real-space scattering-density profile from experimental data. a,b**) Two components ("Phos" and glyerol/carbonyl, "CG") are shown in the headgroup region. Profiles were generated with the SDP program (1). **c,d)** Comparison of EDPs for experimental (Exp) and simulated (Sim) of pure DMPG (c) and pure DPPG (d) lipid bilayers.

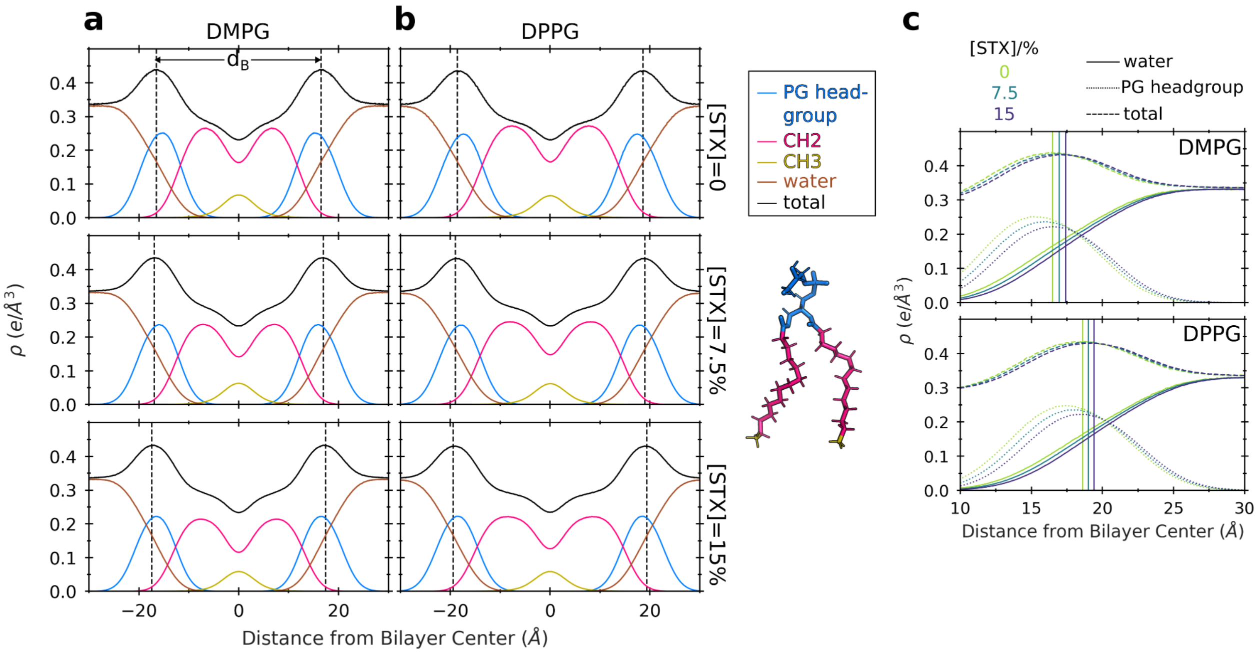


**Figure S4. Electron density profiles of PG component groups, water and the whole system. (a–b)** Profiles were recovered from MD simulations of STX–PG lipid bilayer mixtures with (a) DMPG and (b) DMPG lipids at the indicated STX concentration (rows). The moieties are colored according to the snapshot at the right. The bilayer thickness, $d_B$, was obtained from the decay in the water density (yellow) at positions indicated with the vertical dashed lines (shown here to guide the eye). Note the reduction in the overall density of the distinct PG group components as the STX concentration increases. **(c)** Zoom of the density profiles of water (solid line), PG headgroups (dotted line) and the total density (dashed line) as a function of the STX concentration (color) for the two simulated PG bulk lipids (panels). To guide the eye, the vertical solid lines depict the water density decay point related to the thickness $d_B$. The profiles demonstrate the increase in thickness upon increase in STX concentration.

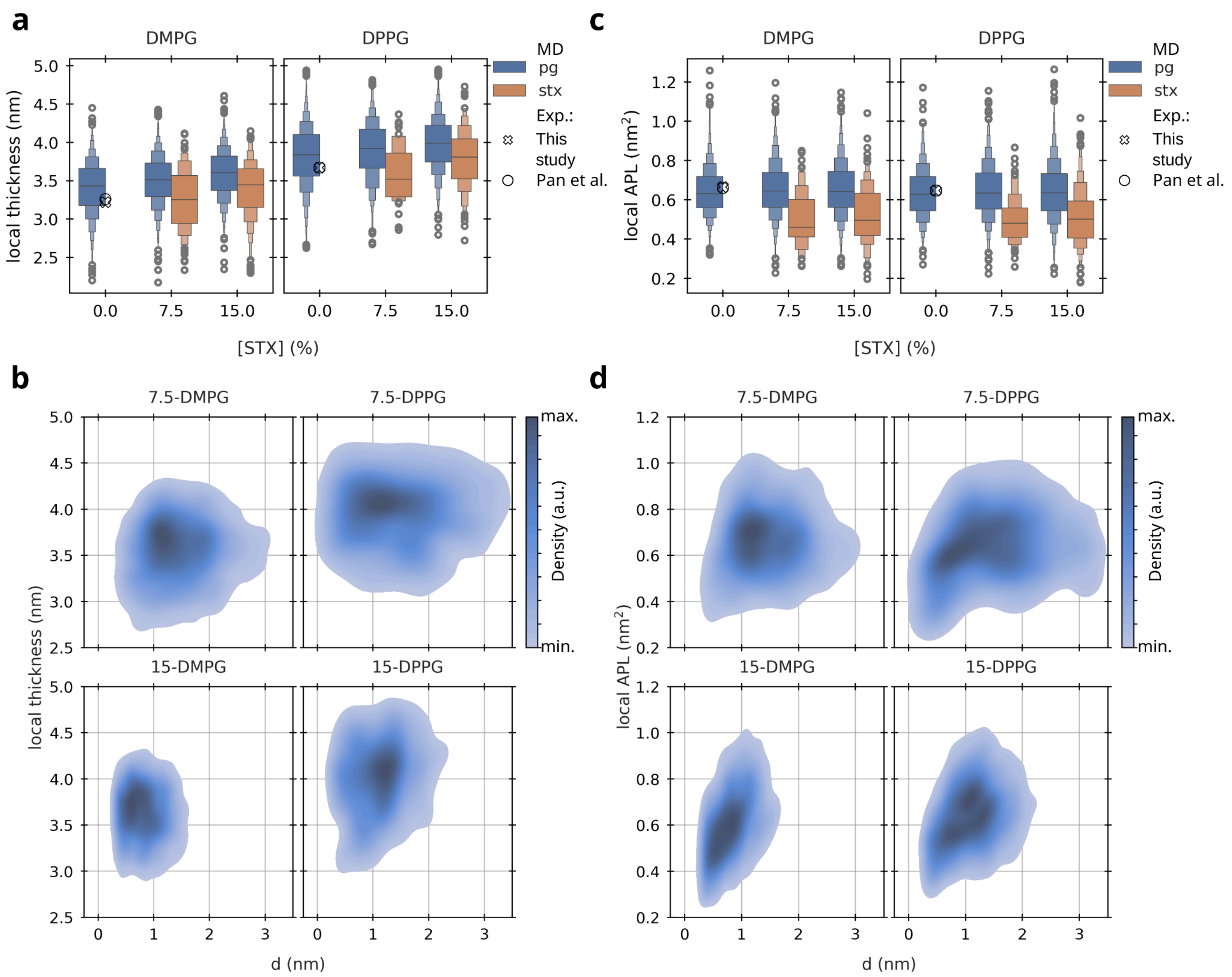


**Figure S5. Local structural details of the thickness (a,b) and area per lipid (APL) (c,d) of STX–PG lipid bilayers recovered from MD simulations. (a,c)** Local thickness (a) and APL (c) distribution is shown separately for PG (blue) and STX (orange), as a function of the STX concentration, for the indicated PG bulk lipids (panels). The distributions are shown as boxen plots. The crosses indicate the experimental values obtained in this study for pure PG bilayers. Previous experimental values from Pan et al. (2) are also shown. **(b,d)** Distribution of the local thickness (b) and local APL (d) for the PG lipids as a function of their distance, d, to the nearest STX molecule. The probability density is represented by color, i.e. the darker the color the higher the probability. The distribution was computed for DMPG lipids (left column) and DPPG lipids (right column) in STX–PG bilayers at 7.5 % mol (top) and 15 % mol (bottom) STX concentrations.

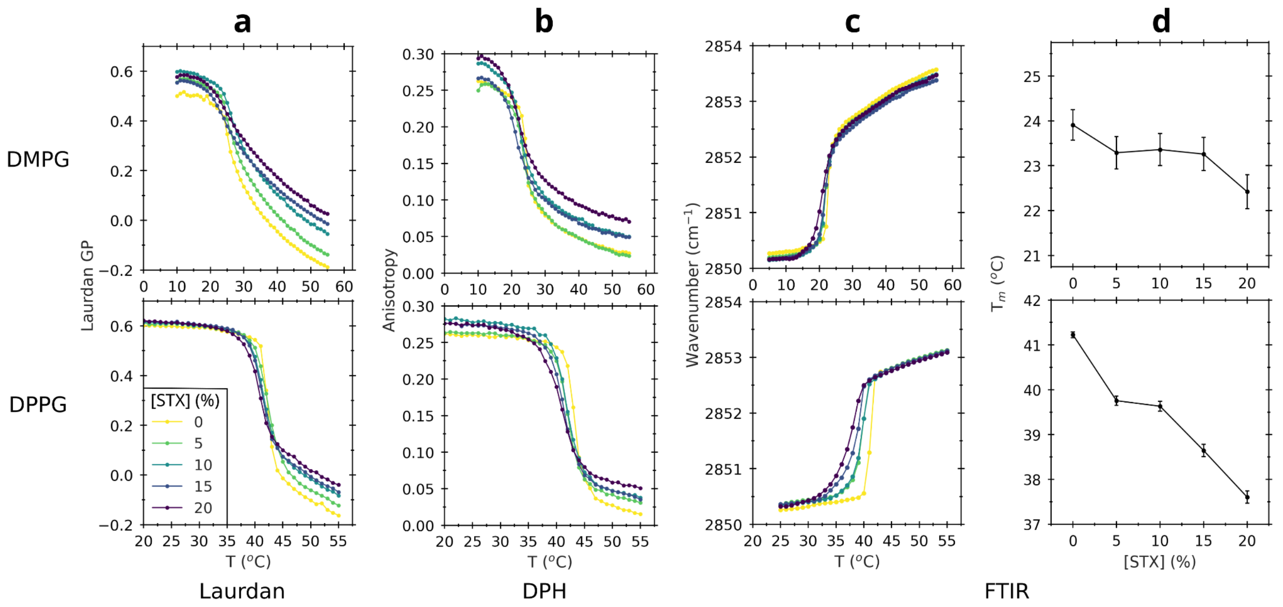


**Figure S6. Staphyloxantin shifts the gel to liquid-crystalline phase transition temperature of DMPG and DPPG lipid bilayers.** Thermotropic curves of DMPG and DPPG membranes with increasing mol% of STX as measured by: **(a)** Laurdan generalized polarization, **(b)** DPH anisotropy, and **(c)** FTIR experiments. In the latter the peak positions of the symmetric stretching vibration band of the methylene groups as a function of temperature were monitored. **d)** Melting temperature, $T_m$, as a function of the STX concentration was obtained by FTIR.

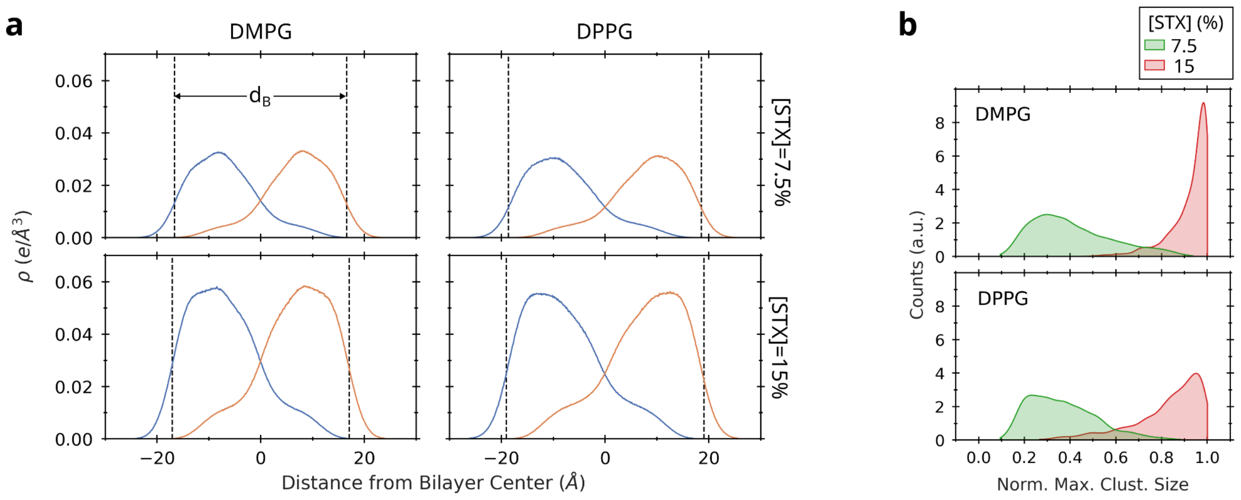


**Figure S7: Interdigitation and clustering of STX lipids. (a)** Electron density profiles of STX as a function of the distance from the bilayer center is depicted separately for each leaflet (blue and orange). The dashed lines indicate the bilayer span, according to the $d_B$ thickness estimate. The density was computed for STX–PG bilayers at the indicated STX concentrations (rows) and bulk PG lipid (columns). **(b)** The maximum cluster size of STX lipids was monitored and distributions are shown for the indicated PG bulk lipids (panels) at the indicated STX concentration (blue and orange). The cluster size was normalized by the STX lipids present in the bilayer (i.e. 32 at 7.5% mol STX and 64 at 15% mol STX concentrations). Accordingly, a value of 1 indicates that all STX lipids were part of a single cluster. The distributions are shown as kernel density estimates.

### References


1. Kučerka, N., J.F. Nagle, J.N. Sachs, S.E. Feller, J. Pencer, A. Jackson, and J. Katsaras. 2008. Lipid Bilayer Structure Determined by the Simultaneous Analysis of Neutron and X-Ray Scattering Data. *Biophys. J.* 95:2356–2367, doi: 10.1529/biophysj.108.132662.

2. Pan, J., D. Marquardt, F.A. Heberle, N. Kučerka, and J. Katsaras. 2014. Revisiting the bilayer structures of fluid phase phosphatidylglycerol lipids: Accounting for exchangeable hydrogens. *Biochim. Biophys. Acta BBA - Biomembr.* 1838:2966–2969, doi: 10.1016/j.bbamem.2014.08.009.
